# Ontogenetic variation in the marine foraging of Atlantic salmon functionally links genomic diversity with a major life history polymorphism

**DOI:** 10.1101/2024.02.26.581711

**Authors:** Tutku Aykanat, Jan Arge Jacobsen, Kjetil Hindar

**Affiliations:** Organismal and Evolutionary Biology Research Program, Faculty of Biological and Environmental Sciences, PO Box 56, FI-00014 University of Helsinki, Finland.; Faroe Marine Research Institute, Nóatún 1, FO-100 Tórshavn, Faroe Islands.; Norwegian Institute for Nature Research (NINA), NO-7485 Trondheim, Norway.

**Keywords:** Feeding strategy, Atlantic salmon, *vgll3*, *six6*, life-history evolution, polymorphism

## Abstract

The ecological role of heritable phenotypic variation in free-living populations remains largely unknown. Knowledge of the genetic basis of functional ecological processes can link genomic and phenotypic diversity, providing insight into polymorphism evolution and how populations respond to environmental changes. By quantifying the marine diet of Atlantic salmon, we assessed how foraging behavior changes along the ontogeny, and in relation to genetic variation in two loci with major effects on age at maturity (*six6* and *vgll3*). We used a two-component, zero-inflated negative binomial model to simultaneously quantify foraging frequency and foraging outcome, separately for fish and crustaceans diets. We found that older salmon forage for both prey types more actively (as evidenced by increased foraging frequency), but with a decreased efficiency (as evidenced by fewer prey in the diet), suggesting an age-dependent shift in foraging dynamics. The *vgll3* locus was linked to age-dependent changes in foraging behavior: younger salmon with *vgll3^LL^* (the genotype associated with late maturation) tended to forage crustaceans more often than those with *vgll3^EE^* (the genotype associated with early maturation), whereas the pattern was reversed in older salmon. *Vgll3 ^LL^* genotype was also linked to a marginal increase in fish acquisition, especially in younger salmon, while *six6* was not a factor explaining the diet variation. Our results suggest a functional role for marine feeding behavior linking genomic diversity at *vgll3* with age at maturity among salmon, with potential age-dependent trade-offs maintaining the genetic variation. A shared genetic basis between dietary ecology and age at maturity likely subjects Atlantic salmon populations to evolution induced by bottom-up changes in marine productivity.

## Introduction

Phenotypic polymorphisms are ubiquitous, discrete trait morphs (e.g., (Bataillon et al., 2022; Funk et al., 2021; Janssen, Bustnes, & Mundy, 2021; Johnston et al., 2013), the genetic basis of which are maintained at higher frequencies than what would be expected under mutation-selection balance (Ford, 1945). Much of the research on phenotypic polymorphism focuses on determining which selective landscapes generate and maintain heritable polymorphisms, such as negative frequency-dependence (Chouteau, Arias, & Joron, 2016; Goldberg et al., 2020; Gross, 1985; Svensson, Willink, Duryea, & Lancaster, 2020), antagonistic pleiotropy (Charmantier, Perrins, McCleery, & Sheldon, 2006; Marden, Langford, Robertson, & Fescemyer, 2021), fluctuating selection (Bonnet & Postma, 2018; Grant & Grant, 2002) and, heterozygote advantage (Coulson et al., 2011; Hedrick, 2012). Furthermore, with the advent of next-generation sequencing, an increasing number of studies have reported associations between discrete chromosomal elements (large-effect loci, inversions, supergenes) and phenotypic polymorphisms (Barson et al., 2015; Funk et al., 2021; Johnston et al., 2013; Matschiner et al., 2022). Such an intense research focus is unsurprising given that studying polymorphisms provides insight into speciation (Jamie & Meier, 2020), one of the cornerstones of biology since Darwin. Despite the increasing number of known genotype-phenotype associations, how such genomic diversity is functionally translated into phenotypic diversity within wild populations is mostly unresolved. This is likely because the experimental studies and sampling designs required for monitoring wild individuals are financially and logistically unfeasible without applying various simplifying assumptions (Mousseau, Sinervo, & Endler, 2000; Reznick, 2016). As such, our understanding of how genomic variation influences individual-level differences in ecology, and how such ecological differences might covary with polymorphic trait states is incomplete.

One well studied genotype-polymorphism association is that between age at maturity and the *vgll3* and *six6* haploblocks in the Atlantic salmon, *Salmo salar* (Ayllon et al., 2015; Barson et al., 2015). Atlantic salmon most often display an anadromous life history; they hatch in rivers where they can spend several years growing before migrating to the ocean. The marine environment is much more productive than freshwater (Gross, Coleman, & McDowall, 1988) and juvenile salmon undertake this costly migration process to maximize growth and lipid storage. This, in turn, supports future reproductive efforts when they return to fresh water to spawn (Esteve, 2005; Ferguson, Reed, Cross, McGinnity, & Prodohl, 2019). Atlantic salmon most commonly stay at sea for one to three years prior to returning to fresh water for reproduction, which is termed as sea age at maturity. Atlantic salmon exhibit a high degree of divergence in this polymorphic phenotype within and among populations, the genetic variation of which is largely explained by two genomic regions (loci), the so-called *vgll3* and *six6*, after the most prominent genes within the haploblock (Barson et al., 2015; Sinclair-Waters et al., 2022). The *vgll3* gene encodes a functionally-conserved transcription co-factor that regulates adiposity in mice (Halperin, Pan, Lusis, & Tontonoz, 2013) and influences pubertal timing in vertebrates (Cousminer et al., 2013). The functional relationship between adiposity and maturation via *vgll3* haploblock likely also operates in Atlantic salmon as condition factor has been shown to drive *vgll3* dependent maturation response in the species (Debes et al., 2021). Salmon with the ‘early’ *EE* genotype are more likely to mature after one year at sea (one-sea-winter salmon, 1SW) than individuals with the ‘late’ *LL* genotype, which are more likely to spend multiple years at sea before maturing (multi-sea-winter, MSW, predominantly composed of 2SW and 3SW salmon). The second major haploblock, *six6*, displays substantial population-level effects on sea-age at maturity among wild salmon populations (Barson et al., 2015), and within aquaculture strains (Sinclair-Waters et al., 2022). The *six6* haploblock contains genes with known functional roles in sensory and digestive system development (Lee et al., 2012; Moustakas-Verho et al., 2020). The molecular functions associated with *vgll3* (adiposity regulation) and *six6* (sensory and digestive system development) suggest that dietary ecology may underpin processes, by which the genomic variation at the *vgll3* and *six6* haploblocks may influence the phenotypic variation in sea-age at maturity.

Variation in how food is acquired by organisms and how derived metabolic energy is invested into different biological processes can be conceptualized using the resource acquisition-allocation framework (Enberg et al., 2012). Briefly, individuals acquire a given amount of a resource from which they service their basal metabolism. Any surplus resources are then allocated to behavior, structural (skeletal/muscle) growth, or maturation/gonadal growth (Enberg et al., 2012). A supposition arising from the resource acquisition-allocation framework is that individuals vary in how much or which resources they acquire, which may alter their maturation trajectories. As such, variation in feeding opportunities and strategies may play a role in the survival and maturation of organisms.

Atlantic salmon is recognized as an opportunistic feeder with extensive ontogenetic and spatiotemporal variation in diet breadth (Rikardsen & Dempson, 2010). Salmon feed increasingly at higher trophic levels as they age, which is strongly correlated to their size (Aykanat et al., 2020; Jacobsen & Hansen, 2001; Salminen, Erkamo, & Salmi, 2001; Salminen, Kuikka, & Erkamo, 1995). The variation in available prey items, as well as preference for particular prey items, can affect the net energy gain from foraging activities, and this may subsequently alter the sea-age at which they mature (Czorlich, Aykanat, Erkinaro, Orell, & Primmer, 2022; Hvidsten et al., 2009; Vollset et al., 2022). For example, Vollset et al. (2022) showed that the reduced marine growth of Atlantic salmon (and subsequent decrease in maturation age) was linked to a reduction in the Arctic cold water flux to the Northeast Atlantic Ocean and potentially to the subsequent decrease in plankton biomass in the feeding grounds in the Northeast Atlantic Ocean. Similarly, using a 40 year time series from a large-sized salmon population, Czorlich et al. (2022) linked the evolutionary response in the *vgll3* genotype towards early maturation to changes in capelin (*Mallotus villosus*) abundance in the Barents Sea, a major prey for Atlantic salmon in the region. In this context, one can posit that genetic variation in ontogeny dependent foraging behavior can be maintained in populations via life-history trade-offs. For example, salmon carrying alleles associated with a foraging behavior that provides higher net-energy gain at early stages at sea will be more likely to acquire enough resources to initiate maturation, and mature at an earlier age (Chaput & Benoit, 2012; Thorpe, Mangel, Metcalfe, & Huntingford, 1998). However, demonstrating the genetic basis for foraging behavior and how this might covary with sea-age at maturity remains highly challenging in salmon due to a lack of adequate samples in the wild. As such, whether foraging behavior varies as a function of genotype, and the evolutionary consequences that this variation could have on the sea-age at maturity phenotype, remains largely unknown. A previous study investigated the age-dependent diet composition as a function of life-history loci in adult Atlantic salmon on their on their return migration along the Norwegian coast (Aykanat et al., 2020). They demonstrated that the *six6* L allele (the allele associated with late maturation) was linked to increased fish prey content, compared to the E allele (the allele associated with early maturation), especially on younger age groups, suggesting a potential role of the locus on maturation norms (Aykanat et al., 2020). However, that study was conducted on Atlantic salmon that had already initiated the maturation process and migration, and with a diet limited to fishes. Therefore, behavioral norms, potential physiological effects of maturation, and the diet-composition do not reflect the conditions during the sub-adult stage in the feeding ground, prior to maturation. Therefore, a shared genetic basis between sea-age and foraging among sub-adult, immature fish (i.e. fish that are still in the oceanic feeding phase, before the onset of maturation) would provide indirect evidence for dietary ecology, acting as a potential process that functionally links genomic diversity at the *vgll3* and *six6* loci with sea-age at maturity in salmon.

An extensive dataset of the diet of wild Atlantic salmon sub-adults (i.e., the marine stage prior to maturation) sampled in the oceanic feeding ground (Jacobsen & Hansen, 2001) provided us with material to test whether ontogenetic variation in foraging behavior is explained by sea-age at maturity loci (*vgll3* – *six6*) in Atlantic salmon. The Atlantic salmon in the study almost exclusively feed on crustaceans and fish, the former of which was more numerous (∼95% of diet items were crustaceans) but comprised only 30% of the content by weight. We propose that a genetic basis in feeding behavior results in variation in net energy gain, which subsequently alters individuals’ probability of reaching the maturation threshold. The salmon diet in our data has a right-skewed distribution with a substantial zero inflation, which we presumed to be shaped by two independent biological processes. The first process explains the “structural-zeros” in the data (i.e., zero inflation), which is used as a proxy to measure the active foraging frequency for particular diet types. The second process is the “count” component, which is used as a proxy to measure the foraging outcome providing that salmon actively pursue prey (see Methods for details). Thus, as a first step, using a zero-inflation model to account for these two processes, we tested whether foraging behavior changes along the ontogeny, and secondly, whether loci associated with sea-age at maturity in salmon, in part, explain the variation in foraging behavior across age groups. Within the same framework, we also estimated whether the variation in diet is determined by age-dependent sex effects, since Atlantic salmon exhibit a sex-dependent maturation pattern (Barson et al., 2015; Fleming & Einum, 2010).

## Methods

### Sampling procedure

The fish samples in this study were originally described in Jacobsen and Hansen (2001). Briefly, Atlantic salmon were sampled from known marine feeding grounds around the Faroe Islands, inhabited by all sea-age groups but predominately by MSW fish (Jacobsen & Hansen, 2001; O’Sullivan et al., 2022). Fish were sampled between November and March (excluding January because of rough sea conditions) during the 1992-’93 and 1993-‘94 fishing periods. Salmon sampled from November-December were considered to have been caught in autumn and those sampled from February-March in winter, which exhibited marked differences in prey diversity, sea surface temperature and stomach content (Jacobsen & Hansen, 2001). The locations of the longlines used to sample the fish were also recorded (*n* = 55, see Fig. S1 in O’Sullivan et al., 2022). Individual salmon had their fork lengths measured, were gutted, and gutted fish were weighed (in kilograms). Gut contents were also removed, identified, and enumerated. Fish scales were taken from each individual salmon for determination of sea-age via scale reading, and for later, DNA extraction for genotyping. The fish sampled in the study were immature, sub-adults with no phenotypic indication of maturation. While early studies indicate as high as 80% of individuals are likely to mature in the following autumn, these estimates are based on limited number of tag recapture studies, and by assessing the maturation probability via hormonal expression using hatchery reared salmon as the baseline (Youngson &, McLay 1984; Jacobsen et al 2001).

However, it is reasonable to assume that virtually all 3SW fish will mature in the following autumn, as 4SW and above sea-ages are rare in salmon populations.

### Dietary quantification and somatic measurements

The stomach content data used in this study is a subset of the data quantified by Jacobsen and Hansen (2001). We did not include data from the 1994-’95 fishing period in Jacobsen and Hansen (2001), as the material for genotyping was not available. The most common diet items among crustaceans were amphipods (e.g., *Themisto* spp.), shrimps (e.g., *Hymenodora glacialis*) and euphausiids (e.g., krill such as *Meganyctiphanes norvegica*), and among fishes were pearlsides (e.g., *Maurolicus muelleri*), lanternfishes (e.g., *Benthosema glaciale* and Myctophidae), barracudinas (e.g., *Notolepis rissoi* and Paralepidae), blue whiting (*Micromesistius poutassou*), and capelin (*Mallotus villosus*) (Supplementary Table 1). Crustaceans were substantially more numerous than fishes (∼95% of diet items were crustaceans), but represented only 30% of the content by weight (Supplementary Table 1). For example, the average size of crustacean species ranged from 0.027 g for *Themisto compressa* to 0.766 for *Sergestes arcticus*, and > 99% were species that were smaller than 0.20 g. The average size of fish subcategories ranged from 0.26 g for fry (mostly consisting of capelin) to 72.0 g for herring, *Clupea harengus* (Supplementary Table 1). For the purposes of this study and to reduce the complexity of data structure and statistical analyses, stomach contents were primarily categorized as crustaceans and fishes.

Apart from the crustaceans and fishes, the diet also contained remains from fish and crustaceans, and organic remains, which were mostly not possible to further identify or enumerate (Supplementary Table 1). In addition, a few stomachs contained bird remains (n=1) and squid (n=11). Fish and crustaceans remains were included in the count-data analysis if they were numerated (n=12 and n=16, respectively). Squids were grouped together with fishes due to their size similarity (excluding them from the analysis did not change any results). One fish with bird remains was excluded from the analysis.

### Genotyping and data filtering

DNA was extracted from the scales of the sampled fish (O’Sullivan et al., 2022). Individuals were genotyped using a GTSeq approach (Campbell, Harmon, & Narum, 2015) with a panel of 167 SNPs as outlined in (Aykanat et al., 2020). This panel allowed individuals to be assigned to their reporting group of origin (O’Sullivan et al., 2022), as well as to score the genotypes at focal SNPs within our target loci. The focal SNPs were *vgll3_TOP_* and *six6_TOP.LD_* (hereafter *vgll3* and *six6*). *Vgll3_TOP_* exhibits the strongest association with sea-age within the haploblock (Barson et al., 2015), while *six6_TOP.LD_* exhibits the second strongest association with sea-age prior to population structure correction (Barson et al., 2015). If an individual was missing a genotype for *vgll3_TOP_*, the missing genotype was inferred from the genotypes of adjacent SNP markers in the same haploblock, which display strong linkage disequilibrium due to close physical proximity to *vgll3_TOP_* (The loci with the strongest linkage with *vgll3_TOP_* was prioritized when imputing the missing genotype, with an order of *vgll3_missense1_*, *vgll3_missens2_*, *akap11*, see, Aykanat, Lindqvist, Pritchard, & Primmer, 2016). Sex was assigned using a combination of phenotypic and genetic sex information, the latter obtained by targeting the sex determination locus (*sdy*, Yano et al., 2013), as outlined in Aykanat et al. (2016). There was 82.3 % concordance between phenotypic and genetic sex, of which we prioritize genetic sexing over phenotypic sexing since sub-adults at sea do not yet develop secondary sexual characteristics, hence phenotypic sex may be more prone to misidentification. Phenotypic sex was used when the genetic sex of an individual could not be determined (n= 39). Individuals lacking genotypic information for either the *vgll3*, *six6* loci, or sex, were excluded from downstream analyses (*n* = 77). A small number of fish with sea-age greater than 3SW were also excluded from the data (n=15). Reporting group was included as a three-level categorical variable, in which an individual was assigned to a reporting group with the highest assignment probability, where the levels corresponded to large-scale geographical regions based on the genetic stock identification in O’Sullivan et al. (2022). These reporting groups were Ireland and the UK (*n* = 473), Southern Norway (*n* = 832), and the Northern group (*n* = 271), the latter consisting of fish from Northern Norway, including the Teno River, Eastern Finnmark/Northern coast of Kola, and the Barents/White Sea reporting groups. In total, 1536 individual salmon consisting of 168 1SW, 950 2SW, and 418 3SW individuals were used in the downstream analyses (Table 1).

**Table 1:** The *vgll3* and *six6* frequencies across sea-ages and sexes in the dataset. Number of individuals in each group are indicated in parenthesis.

| Sex | Sea-age | *vgll3* |  |  | *six6* |  |  |
| --- | --- | --- | --- | --- | --- | --- | --- |
|  |  | EE | EL | LL | EE | EL | LL |
| Female | 1SW | 0.56 (49) | 0.25 (22) | 0.19 (17) | 0.42 (37) | 0.32 (28) | 0.26 (23) |
|  | 2SW | 0.33 (216) | 0.44 (293) | 0.23 (153) | 0.15 (99) | 0.34 (226) | 0.51 (337) |
|  | 3SW | 0.11 (30) | 0.38 (108) | 0.51 (144) | 0.02 (6) | 0.21 (58) | 0.77 (218) |
| Male | 1SW | 0.43 (34) | 0.41 (33) | 0.16 (13) | 0.30 (24) | 0.43 (34) | 0.28 (22) |
|  | 2SW | 0.26 (74) | 0.31 (89) | 0.43 (125) | 0.13 (36) | 0.29 (83) | 0.59 (169) |
|  | 3SW | 0.04 (5) | 0.21 (29) | 0.75 (102) | 0.01 (2) | 0.20 (27) | 0.79 (107) |

### Statistical analyses

We modelled the total number of crustaceans and the total number of fishes in the stomachs of individual salmon as a pair of separate two-component zero-inflation models, for which 46.1 and 79.9 % of stomachs had no crustaceans and fishes, respectively (Supplementary Figure 1, Supplementary Table 2). Zero-inflation models simultaneously estimate two independent processes that shapes the distribution of the response variable (diet in the stomach), which has been suggested for overdispersed and zero-inflated data (Blasco-Moreno et al., 2019). The first process was modelled with the zero inflation component (using binomial error distribution), and quantified for the “structural-zeros” (i.e., inflated zero counts) in the data that were due to individuals not actively foraging for the specific prey type. We used this statistic as a proxy to measure active foraging frequency. The second process was modelled with the count component (using a negative binomial error distribution that includes the “random zeroes” in the data) that were due to sampling variation when salmon actively forage for prey. We used this statistic as a proxy to measure the outcome of active feeding, i.e., foraging outcome. We modelled the data using a negative binomial error distribution, rather than Poisson, because the data were overdispersed (residual mean and variance were substantially unequal) violating the main assumption of the Poisson error distribution (Blasco-Moreno et al., 2019). Similarly, we did not use a truncated model (i.e., zero altered model), which assumes that all the zeroes in the data originates from one process (i.e., no active foraging), while some of the zeroes in the data might be due to random failure during active foraging. As such, the inflated model can simultaneously account for two different statistical types of zeroes (Blasco-Moreno et al., 2019; Martin et al., 2005).

The model structure (for both components) is as follows:

Y = μ + Season + fishing period + Reporting Group + Sea-Age x (Sex + *vgll3* + *six6* + *L_age.std_)* + ε*_location_* + ε_error_

whereby, for both crustacean and fish models, season (autumn, winter), fishing period (1992-‘93, 1993-‘94), and reporting group were included as categorical fixed effects. The *vgll3* and *six6* genotypes, as well as sex and age-standardized length (*L_age.std_*) were modelled by interacting with sea-age, since we quantified a sea-age-dependent diet response associated with these variables. As expected, the genetic variation in both *vgll3* and *six6* loci was substantially linked to sea-age in both sexes, likely due to increasing representation of the late maturing allele in older age groups (Table 1), but this does not introduce any bias when modeling diet since the genotype is modelled by interacting with sea-age. All covariates interacting with sea-age (*vgll3*, *six6, sex, and L_age.std_*) were scaled by their respective means and standard deviations to improve the convergence, and interpretability of the coefficient tables. Spatial variation in sampling effort (sampling location, N=55) was included in the model as a random effect.

To evaluate whether any taxonomic groups within crustaceans drive the relationship between genotypes and diet, we further broke down the response variable into lower order taxonomic groups when numbers within specific taxa were sufficiently large. For that, we reanalyzed the data using Hyperiid amphipods (*Themisto* spp.) and Euphausiids as response variables, which comprised the majority of the crustacean diet in the stomach (Supplementary Table 1). The representative sizes of the diet between these two groups were also different, with the latter being substantially larger than the former (Supplementary Table 1). We did not implement the same approach for fishes that had restrictively small number of prey items per species.

The unequal contribution of prey taxa with different digestion rates (and overestimation of slowly digested prey items) is an inherent problem in stomach content analyses (Amundsen and Sánchez-Hernández, 2019). Yet, the main contrasts of interest in the study (i.e. genetic variation in *vgll3* and *six6*) was quantified independently within different diet types, and our results should be robust to such effects, assuming that the genetic variation in these loci is not associated with differential digestion rate.

All models were run using the *glmmTMB* package (version 1.1.5, Brooks et al., 2017) in R version 4.1.0 (R Core Team, 2019), using a negative binomial error structure with a logit link function for the count component, and with a binomial error structure with a logit link function for the zero-inflation (zi) component, using default optimizer settings. Marginal means for genotype effects and their confidence intervals were calculated using the *emmeans* package (version, 1.8.2, Lenth, 2023). Model diagnostics were carried out using *DHARMa* package in R (version 0.4.5, Hartig 2022), whereby randomized quantile residuals were simulated for both models to determine if any explanatory covariate displayed heteroscedasticity (Hartig 2022). Using the aforementioned two-component models, we primarily asked whether sub-adult foraging behavior varies as a function of life history genotype and age. We also tested whether sex had an age-dependent effect on adult foraging behavior.

## Results

### Crustaceans model

*Sea-age:* Based on the zero-inflation models, older sea-age groups were more likely to forage for crustaceans (zi component: z = -3.196, *P* = 0.001), but younger sea-age groups had substantially higher number of crustaceans in the stomach, indicating a higher foraging outcome (count component: z= -4.478, *P* < 0.001, Supplementary Table 3, Figure 1).

**Figure 1:**
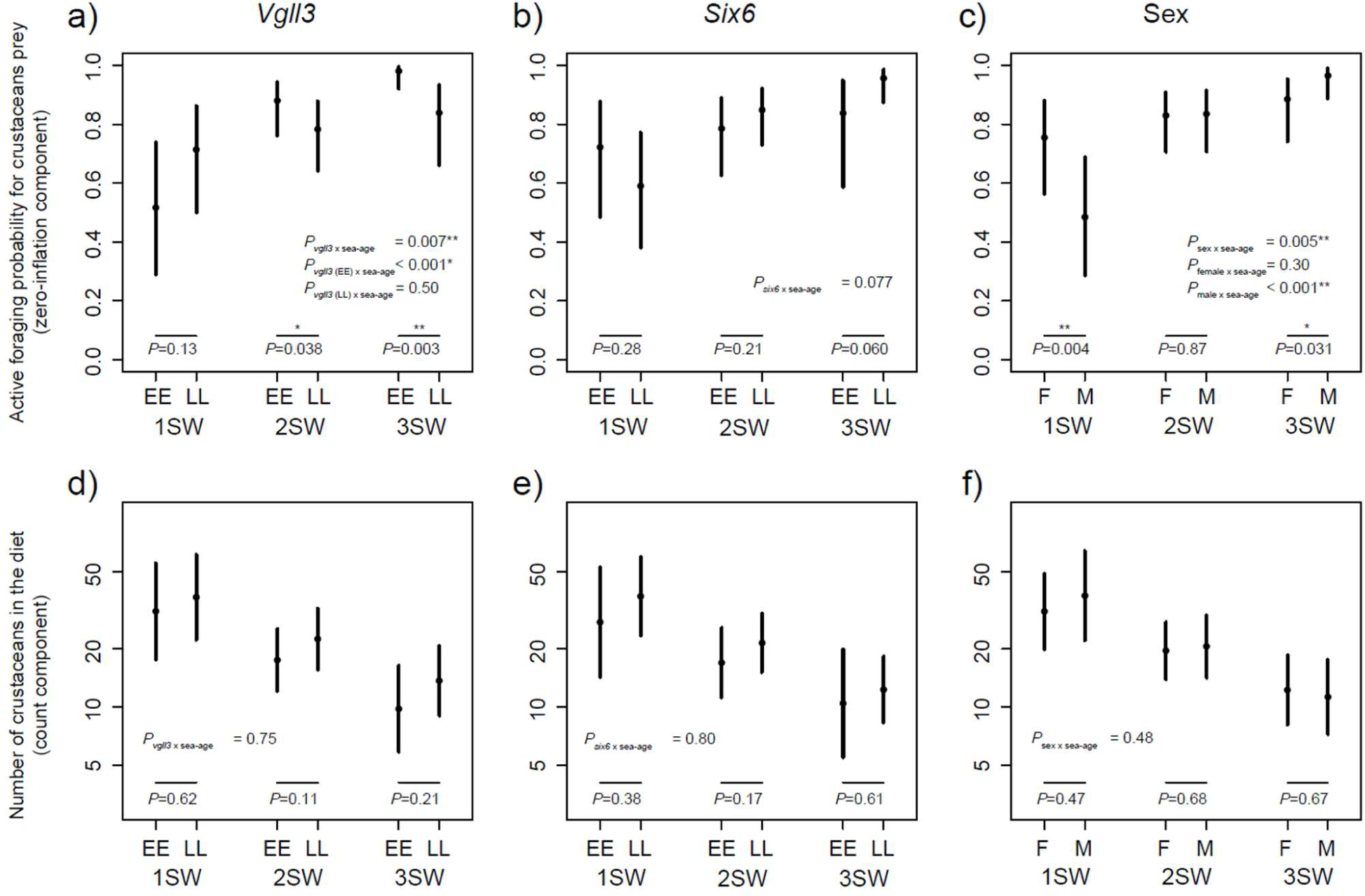
Crustaceans acquisition patterns of sub-adult Atlantic salmon as a function of sea-age-dependent genotype (*vgll3* and *six6*) and sex. Upper and lower panels indicate marginal means for foraging frequency (zero-inflation component), and foraging outcome (count component), respectively. Error bars are 95% confidence intervals. (The y-axes in d,e, and f are log-scaled. Asterisks indicate *< 0.05 and, **< 0.01.)

#### Vgll3 and six6

Genetic variation in the *vgll3* locus was strongly associated with an age-dependent change in crustacean foraging frequency (zi component: z = 2.690, *P*_sea-age x *vgll3*_ = 0.007, Figure 1a, Supplementary Table 3). As such, *vgll3*^LL^ genotype was insignificantly associated with increased foraging in younger sea-age groups (Odds ratio = 2.33, 95% CI = 0.78 – 7.04, *P* = 0.132, Figure 1d), but in the older sea-age groups, the alternative, *vgll3*^EE^ genotype, was significantly associated with increased foraging for crustaceans (for 2SW, odds ratio = 2.04, 95% CI = 1.04– 3.98, *P* = 0.038; for 3SW, odds ratio = 9.67, 95% CI = 2.15– 43.57, *P* = 0.003, Figure 1a). The difference appeared to be driven by the *vgll3*^E^ allele, which exerts a significant age-dependent response to crustacean’s foraging frequency in the diet (z = -4.081, *P*_sea-age|*vgll3EE*_ < 0.001), whereas the *vgll3*^L^ allele had no age-dependent effect (z = - 1.112, *P*_sea-age|*vgll3EE*_ = 0.507, Figure 1a). However, *vgll3* genotypes did not alter the foraging outcome, as there was no difference in the abundance of crustaceans in salmon diet in relation to *vgll3* genotype (Figure 1d).

Crustacean diet as a function of *six6* locus was insignificant for both foraging frequency (zero-inflation components) and foraging outcome (zero-inflation components, Figure 1b,e, Supplementary Table 3).

#### Sex

Sex exhibited a strong age-dependent association with foraging frequency (zi component: z = -2.822, *P* = 0.005, Figure 1c), but did not alter the foraging outcome (count component: z = -0.703, *P* = 0.482, Figure 1f). Female salmon foraged for crustacean more often than males in the 1SW group (Odds ratio = 3.27, 95% CI = 1.46 – 7.31, *P* = 0.0042), whereas the pattern reversed in the older sea-age groups and the 3SW males foraged crustaceans more often than the 3SW females (Odds ratio = 3.55, 95% CI = 1.13 – 11.21, *P* = 0.031, Figure 1c). The difference appeared to be driven by a greater age-dependent response by males (z = -4.297, *P*_sea-age x females_ < 0.001), compared to females (z = -1.489, *P*_sea-age x males_ = 0.296, Figure 1c).

#### Season, year, reporting group and length effects

Season was an important determinant for crustacean diet. Salmon were more likely to forage for crustaceans in the winter (zi component: *z* = - 2.439, *P* = 0.015), but had an overall lower abundance of prey in the diet (count component: *z =* -5.107, *P* < 0.001, Supplementary Table 3). Fishing period did not have a significant effect on foraging frequency, yet the number of crustaceans in the diet was significantly different between the two sampling years (count component: *z =* 5.512, *P* < 0.001, Supplementary Table 3).

Reporting group generally did not affect the variation in the crustacean diet, but fish assigned to the Northern Norway reporting group had marginally fewer crustaceans in the diet than the fish assigned to Ireland and the UK reporting group (*z* = -2.504, *P* = 0.037, Supplementary Table 3). Age-standardized length as a function of sea-age (*L*_age.std_) was not a significant factor for the crustaceans diet (Supplementary Table 3). Sequential exclusion of these variables from the model (season, fishing period, reporting groups, and *L*_age.std_) did not meaningfully influence the genotype or sex effects observed in the crustacean diet (data not shown).

#### Alternative parameterization of variables

When the crustacean diet was subdivided into lower taxonomic orders, two major groups (Hyperiid amphipods and Euphausiids) appeared to exhibit similar effects and effect sizes (and similar to the overall crustaceans model) associated with the *vgll3* locus, albeit not always significant due to lower sample sizes (Supplementary Table 4 and 5), suggesting that neither a particular group nor size drives the age-dependent genetic or sex relation.

In the implemented framework, we modelled the life-history loci and sea-age as a continuous variable, assuming additivity for the latter, and linear relation of the response variable with the former. This model structure was substantially more parsimonious than models that assume non-additivity (where genotypes were modelled as categorical variables), and/or non-linear age-dependent response (where sea-age was modelled as a categorical variable, Supplementary Table 6). Similarly, a model that accounts for different combinations of three way interaction between sea-age, sex, and genotype, which was previously suggested as a mechanism for a sex dependent genetic effect on sea-age at maturity (i.e., for *vgll3*, Barson et al., 2015), was not supported (Supplementary Table 7).

#### Convergence and diagnostics

Convergence was achieved in all models, enabling us to compare different parameterizations of sea-age and genotype. Overall, the diagnostics for the crustaceans model suggested a good model fit (Supplementary Figure 2a).

### Fish model

#### Sea-age

Similar to the crustaceans model, older sea-age groups were more likely to forage for fish (zi component: z = -3.704, *P <* 0.001), and younger sea-age groups exhibited higher foraging outcome (count component: z= -4.204, *P* < 0.001, Supplementary Table 8, Figure 2).

**Figure 2:**
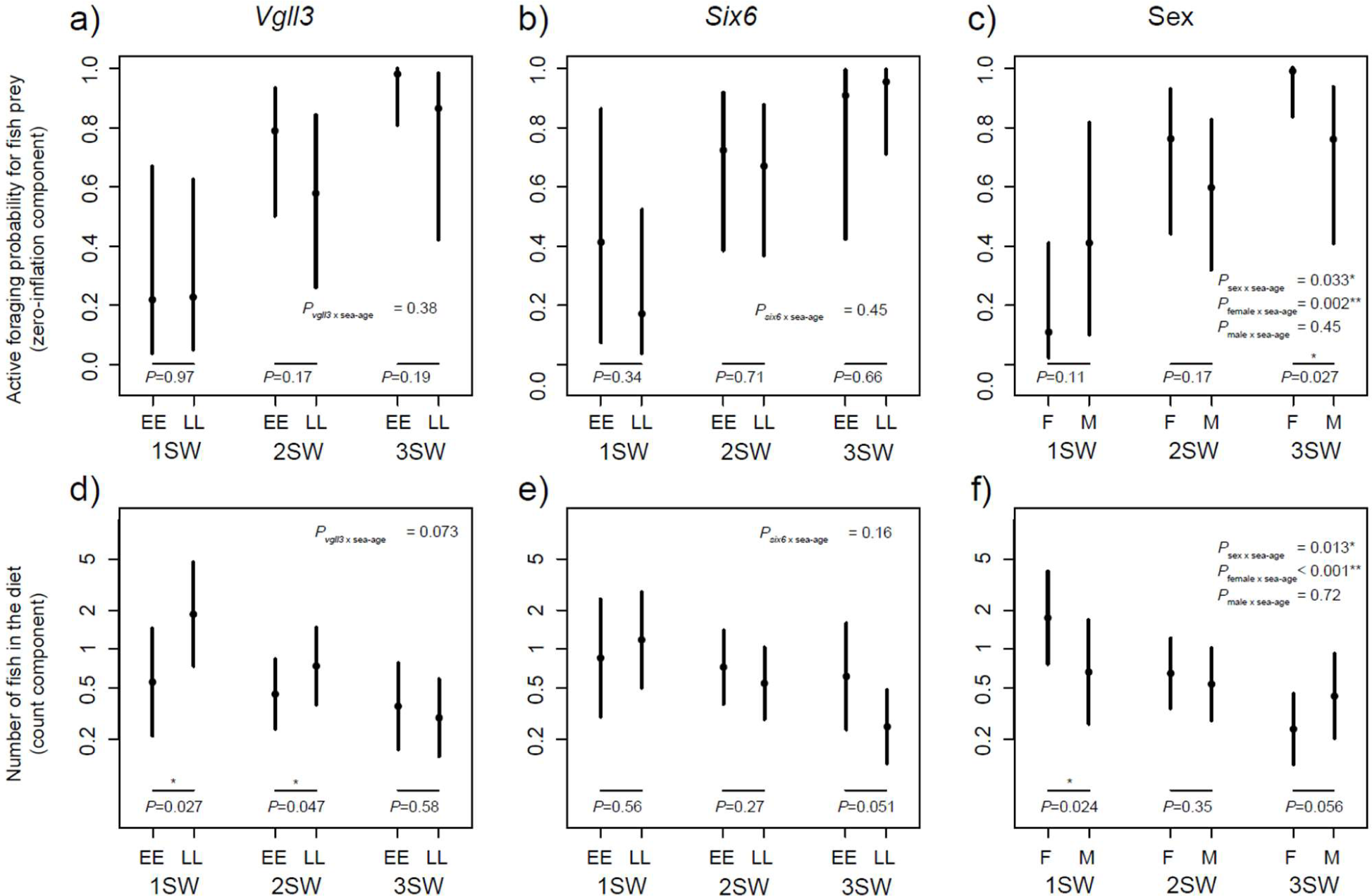
Fish acquisition patterns of immature Atlantic salmon as a function of sea-age-dependent genotype (*vgll3* and *six6*) and sex. Upper and lower panels indicate marginal means for foraging frequency (zero-inflation component), and foraging outcome (count component), respectively. Error bars are 95% confidence intervals. (The y-axes in d,e, and f are log-scaled. Asterisks indicate *< 0.05 and, **< 0.01.)

#### Vgll3 and six6

Frequency of foraging for fish was generally insignificant for *vgll3* and *six6* loci. The foraging outcome, however, was significantly higher for *vgll3*^L^ allele (count component: z = 2.088, *P* = 0.037, Supplementary Table 8), whereby the effect was diminished in older age-groups due to a marginal age-dependent effect (count component: z = -1.796, *P*_sea-age x *vgll3*_ = 0.073, Supplementary Table 8). Salmon with *vgll3*^LL^ genotype had more fish in the stomach than salmon with *vgll3*^EE^ genotype in 1SW and 2SW sea-age groups (for 1SW, odds ratio = 3.36, 95% CI = 1.15 – 9.81, *P* = 0.027; for 2SW, odds ratio = 1.65, 95% CI = 1.01 – 2.71, *P* = 0.047), but the difference was negligible in the 3SW age group (odds ratio = 0.81, 95% CI = 0.39 – 1.70, *P* = 0.582, Figure 2d). *Six6* was not a significant factor affecting foraging outcome in the fish diet.

#### Sex

Sex had a marginal effect on foraging frequency and outcome for fish diet. Frequency of foraging for fish increased in females as salmon gets older (zi component: z = 2.127, *P*_sea-age x *sex*_ = 0.033, Figure 2c, Supplementary Table 8), and females foraged for fish more often than males in the 3SW age group (odds ratio = 26.506, 95% CI = 1.45 – 483.96, *P* = 0.027, Figure 2c). In contrast, the number of fish in the diet was higher in females in 1SW age groups (odds ratio = 2.64, 95% CI = 1.14 – 6.14, *P* = 0.024, Figure 2f, Supplementary Table 5), and the pattern was reversed in 3SW, and males had marginally higher fish diet in the 3SW age group (odds ratio = 1.80, 95% CI = 0.99 – 3.29, *P* = 0.055, Figure 2f) suggesting an age specific sex pattern in foraging outcome (count component: z = 2.484, *P*_sea-age x *sex*_ = 0.013). For both foraging frequency and outcome the age-dependent effects were driven by the age-dependent response of females, rather than males (Figure 2c, f).

#### Season, year, reporting group and length effects

Season was an important determinant of the fish diet. Salmon foraged for fish more often in winter (zi component: *z* = -2.014, *P* = 0.044), and acquired more fish (count component: *z =* 4.413, *P* < 0.001, Supplementary Table 8). Fishing period, reporting group, or age standardized length did not have a significant effect on the fish diet (Supplementary Table 8).

The estimated effects of season or reporting group did not meaningfully influence the genotype or sex effects observed, as the sequential exclusion of these parameters did not meaningfully change the effect size and/or significance (data not shown). However, the model did not converge when fishing period was excluded from the model, therefore their effect on the focal variables was not assessed. On the other hand, the marginal *vgll3* effect on foraging outcome appeared to improve and became significant after excluding *L*_age.std_ from the model, suggesting possible covariation between the two (Supplementary Figure 3). As such, salmon with *vgll3*^LL^ genotype had more fish in the stomach than salmon with *vgll3*^EE^ genotype in 1SW and 2SW sea-age groups (for 1SW, odds ratio = 4.64, 95% CI = 1.52 – 14.16, *P* = 0.007; for 2SW, odds ratio = 2.02, 95% CI = 1.22 – 3.35, *P* = 0.006), but the difference was negligible in the 3SW age group (odds ratio = 0.88, 95% CI = 0.41 – 1.90, *P* = 0.746, Supplementary Figure 3).

#### Alternative parameterization of variables

For the fish diet model, the *vgll3* and *six6* genotypes were modelled additively (EE, EL, and LL genotypes were coded numerically by 1, 2, and 3) which were more parsimonious than non-additive models (i.e., when genotype values were categorically modelled. Supplementary Table 6.). Sea-age was also parameterized as a continuous variable, but comparing continuous and categorical parameterization was not possible due to convergence problems with the latter (Supplementary Table 6). Similar to the crustaceans model, in the fish model, a model that accounts for different combinations of three-way interaction between sea-age, sex, and genotype was not supported, although we did not achieve convergence in the four-way interaction model that includes sea-age, sex, and two loci of interest (Supplementary Table 7).

#### Convergence and diagnostics

As evidenced by the diagnostics plot obtained from the DHARMa package, the fish model had suboptimal fitting (Supplementary Figure 2b). Yet, this shortcoming may have had little impact on the main conclusions. For example, when additional dispersion parameters were added to account for variance heterogeneity for fishing season, reporting group, season and sex, the homogeneity of variance (or within group deviation from uniformity) was achieved for all parameters except for fishing season, and the Kolmogorov–Smirnov deviation test was not significant anymore (Supplementary Figure 2c). The addition of these dispersion parameters did not alter the conclusions associated with genetic or sex effects (Supplementary Table 9). Convergence was not achieved when testing for alternative parameterization of model covariates, such as when sea-age and genotype effects were modelled as a categorical variables, or when assessing the effect of non-focal variables on the main conclusions (by sequentially excluding them from the model). Therefore, the model results from the continuous parametrization of sea-age should be approached cautiously.

## Discussion

We demonstrated an age-dependent genetic pattern in marine diet variation in wild, sub-adult Atlantic salmon. By simultaneously capturing the effects of two biological processes that shape diet variation, namely active foraging frequency (via the zero-inflated component) and foraging outcome (via the count component), we provided insights into the foraging dynamics of Atlantic salmon at sea.

Intriguingly, the two processes had remarkably different sea-age-dependent patterns: younger individuals contained more crustaceans in the stomach than older age groups suggesting a higher foraging outcome, despite lower foraging frequencies. This pattern cannot be explained by the higher assimilation rate of older salmon (i.e. those with larger guts) as higher assimilation rates should also correlate with higher frequencies of empty stomach. The results from these two biological processes combined, suggest that older sub-adult salmon adopts a continuous feeding strategy, albeit a less efficient one, perhaps to fulfill the energetic demands of a larger body size (Jonsson & Jonsson, 2011; Nina Jonsson & Jonsson, 2003). On the other hand, these results are in contrast with previous findings by (Aykanat et al., 2020) which quantified the diet of mature Atlantic salmon in coastal waters, on their migration to freshwater for reproduction. Their study showed that older salmon exhibited a higher frequency of empty stomach, but had a heavier diet content when the stomach was non-empty, an indication of a feast and famine strategy as a foraging adaptation to maintain a high energy balance. Likewise, the disparity in resource acquisition patterns was also evident in the ontogenetic shift in diet, with the former study exhibiting a clear ontogenetic shift in diet towards larger fish species in older salmon, whereas in the current study, older salmon acquired fewer diet items irrespective of taxa (crustaceans *vs.* fish, Supplementary Table 3 and 8). It is unclear what underlies these opposing patterns between these two studies, but the stark differences in the environments (coastal *vs.* oceanic feeding grounds with substantial differences in diet composition) and life stages of salmon (immature sub-adults *vs.* mature adults) appear to alter feeding strategies. For example, the diet was primarily composed of four fish species in Aykanat et al. (2020), a substantial portion of which was larger than three grams while in the present study only a small number of fish species in the diet had, on average, weighed more than three grams, indicating a striking difference in diet breadth. Combined, these findings suggest that resource acquisition strategies are strongly dependent on the life stages at sea, and that conclusions cannot be generalized across life stages.

Our results indicate that older sea-age groups feed more frequently despite that they lack the foraging outcome of younger age groups. This is especially intriguing in relation to the crustaceans diet, as it is contrary to the expected patterns of ontogenetic diet shifts in salmon (Rikardsen & Dempson, 2010). The larger body of older sea-age groups is energetically more demanding (Jonsson & Jonsson, 2011; Jonsson & Jonsson, 2003), and it might be that older salmon are in need of constant searching for food to support the energetic demands of larger body, relative to younger age groups, especially when prey size is small, a strategy of which may come in the expense of efficiency of foraging. It could also be speculated that salmon partly use filter feeding using their gill-rakers as sieve during high density crustacean prey distributions. Small salmon (parr and smolts) use filter feeding to catch small prey (e.g. Keeley & Grant, 1997). If ocean feeding post-smolts and small salmon also use this technique, acknowledging that gill-raker spacing is proportional to fish length (Wankowski 1979), 1SW salmon might benefit from their smaller gill-raker spacing when targeting relatively small crustacean prey compared to larger MSW salmon.

We also demonstrate a shared genetic basis between a major life-history polymorphism (sea-age) and feeding strategy in Atlantic salmon with substantial sea-age-dependence. Specifically, the age-dependent patterns associated with the *vgll3* locus, predominantly in foraging rates of crustaceans, provide evidence that feeding strategy is a functional ecological process by which differences between sea-age phenotypes might be realized. This is a rare example of genetically-controlled differences in resource acquisition of a highly migratory fish species, and demonstrates that salmon could be subject to selection if oceanic food webs change (Czorlich et al., 2022; Strøm, Ugedal, Rikardsen, & Thorstad, 2023; Vollset et al., 2022). Crustaceans are a primary source of dietary material for Atlantic salmon in the Northeastern Atlantic Ocean (Jacobsen & Hansen, 2001; Rikardsen & Dempson, 2010). A reduction in the water flux from the colder Arctic region into the northern Atlantic around 2005 was associated with a decrease in biotic and abiotic factors including plankton abundance, as well as the specific growth rate of salmon, the latter which may eventually result in a later maturation age (Vollset et al., 2022). While our results do not support the idea that *vgll3* is linked to a change in feeding efficacy (as quantified by the count component of the model), foraging for crustaceans (as quantified by the zi component) appeared to be regulated by the *vgll3* locus. These results suggest that an ecological shift in the northeast Atlantic Ocean might exert an evolutionary change in the *vgll3* locus.

Crustacean acquisition patterns in relation to *vgll3* might provide some clues on how the variation in sea-age at maturity may be mediated via a genetic polymorphism in resource acquisition. To illustrate, 1SW salmon with the *vgll3*^*EE^ genotype appear to obtain as many crustaceans as 1SW salmon with the *vgll3\**^LL^ genotype (Figure 1d), despite foraging for crustaceans less frequently (Figure 1a). In other words, 1SW individuals with *vgll3*^EE^ genotype gain a similar amount of food (estimated from count model) with less active foraging. Such an increase in acquisition efficiency may be one of the factors that allows salmon with the *vgll3*^EE^ genotype to mature earlier as 1SW fish, i.e., by allocating surplus resources (saved through efficient foraging) towards fat deposition and reproductive investment. Furthermore, we also observed a parallelism in age-dependent crustaceans foraging activity between *vgll3* genotypes and sex, and in relation to their effect on maturation timing. For example, females, which mature later than males (Barson et al., 2015; Fleming & Einum, 2010), forage for crustaceans more often than males (and with similar acquisition efficiencies), a pattern that is similar between *vgll3^LL^* and *vgll3*^EE^ genotypes. Similar to salmon with *vgll3*^LL^ genotype, perhaps, foraging behavior of females may not be sufficient to acquire energy to trigger early maturation. An optimal foraging strategy for earlier maturation might be to decrease active foraging time and increase foraging efficiency. On the other hand, it is not clear whether age-dependent changes in diet patterns associated with early maturation in the *vgll3*^EE^ genotype and males could indicate an age-dependent trade-off associated with early maturation. In fact, age-dependent changes in foraging time were mainly driven by age-dependent changes in the *vgll3*^EE^ genotype, and in males, both of which are associated with early maturation. Based on the foraging patterns, the early maturation genotype and males are becoming less efficient in acquiring crustaceans in older sea-ages (i.e., 3SW). Thus, we propose that early maturation genotype and males have evolved to have higher foraging efficiencies for crustaceans in early ages, potentially linked to earlier maturation, but this may come with an efficiency trade-off in later years at sea. In turn, a seemingly maladapted foraging strategy at younger ages might be selected for in salmon populations by eventual delay in maturation timing, resulting in maturation at a larger size.

The crustacean acquisition model provides a functional ecological process explaining genetic (via the *vgll3* locus) and sexual basis of maturation timing. However, it is not clear if this effect is manifested via condition factor, a primary phenotypic trait associated with maturation timing (Thorpe et al., 1998) which is linked to *vgll3* during precocious male maturation (Debes et al., 2021). Using the same dataset, Aykanat et al. (2024) showed an ontogenetic trade-off between condition factor and diet composition, whereby an increased crustaceans diet increases the condition factor in younger sea-age groups but decreases it in the older age groups, while fish diet exhibited the opposite effect. To explore whether *vgll3* or sex was linked to age-dependent differences in the condition factor, and consistently so with the expected maturation patterns, we implemented a linear mixed model with a Gaussian error structure using condition factor as the response variable, and using a similar covariate structure as above (except that sea-age was modeled categorically which was more parsimonious than a numeric model). There was no evidence of an age-dependent association between the condition factor and *vgll3* structure (Supplementary Table 10). Males, on the other hand, exhibited a higher condition factor in the 1SW sea-age group (z = 1.982, *P* = 0.0476) and a lower condition factor in 2SW (z = -4.776, *P* < 0.001) compared to females, but not in the 3SW sea-age group (t = -0.282, *P* = 0.7779). This is consistent with the expected maturation patterns in Atlantic salmon, where 1SW sea-age group is more predominately males, in contrast to proportionally more females in the 2SW age-group (Fleming & Einum, 2010), suggesting that sex specific diet differences might drive maturation via the effect of condition factor. However, the condition factor used here was calculated using the gutted weight, hence it did not include any variation in condition that might be linked to the guts and visceral tissue, both of which are likely important ecological traits in the processes of maturation (Mobley et al., 2021).

Unlike crustaceans, fish acquisition was generally similar between genotypes across sea-age groups. However, foraging outcome was meaningfully different within *vgll3* genotypes and sexes across sea-ages, whereby *vgll3*^LL^ genotype and females consumed a higher number of fish in the 1SW age group with marginal and significant age-dependent changes for *vgll3* and sex, respectively. This pattern is concordant with the notion that fish acquisition may not be an efficient energy source for 1SW for early maturation (Aykanat et al. 2024), and further highlights resource acquisition as a potential ecological function that links maturation timing to genetic polymorphisms in the *vgll3* locus, and sex. However, caution is required when interpreting the results for the fish model, as the model fit deviates from the linear model assumptions and convergence for alternative parameterization options was not achieved.

### Caveats

Studying the evolutionary ecology of Atlantic salmon in the marine phase provides unprecedented possibilities for understanding the ecological and evolutionary basis of life history variation in the wild, but is inherently difficult. This is due to their extensive oceanic migration with substantial spatio-temporal variation both within (Gilbey et al., 2021) and between age-classes. Such stochasticity (due to the marine ecology of salmon and the sampling methods) results in structurally complicated data. It was impossible to quantify and compare all possible model structures to account for this complexity. The complex model structure also resulted in convergence issues, especially with alternative parameterization of the fish model, which limited the exploration of potential confounding effects. However, especially for the crustaceans model, we were able to assess the effect of alternative parameterizations to a great extent, which appears to not affect the main conclusions.

1SW salmon utilize the feeding grounds around the Faroe Islands relatively less often than older age groups (O’Sullivan et al., 2022), suggesting that 1SW fish have spatially distinct feeding ranges in the ocean. Therefore, it is unclear whether the patterns in this study were also observed on feeding grounds that are predominantly used by 1SW salmon. Likewise, the samples were acquired in the early 90s, about three decades prior to the present date. Since then, especially in the 2000s, the Northeast Atlantic Ocean underwent a large-scale regime shift with a substantial increase in temperature, and a decrease in planktonic biomass likely due to large-scale changes in the proportion of cold Arctic water flowing to the regions, followed by changes in the demographic structure Atlantic salmon towards a higher proportion of older age groups (Vollset et al., 2022). It is unclear how these changes in the environment would impact the genetic-diet associations observed in this study. Combined, this encourages further work to verify potential acquisition patterns across time and space, and to better understand acquisition-dependent life history variation associated with the *vgll3* haploblock. Similarly, the low numbers in 1SW are associated with large 95% prediction intervals around 1SW. This limits the inferential strength of the relevant results, and predictive power of the analyses.

Given the complex data structure and associated challenges with the interpretations, we limited our analyses to asking broad questions concerning the eco-evolutionary dynamics of marine feeding strategies and sea-age phenotype. As such, we primarily grouped diet into two major categories (crustaceans and fish), and did not address more detailed or refined hypotheses, due to a lack of statistical power to do so robustly. However, two major crustacean taxa (Hyperiid amphipods and Euphausiids) had sufficient data to achieve model convergence, by which we concluded that the general patterns observed in crustaceans were similarly reflected in these taxa, also irrespective of their size distribution, suggesting that prey size may not be important in shaping age-dependent genetic and sex diversity on diet acquisition.

### Conclusions

An important consequence of anthropogenic change has been the large-scale disturbance of ecosystems globally (Sydeman, Poloczanska, Reed, & Thompson, 2015; Vollset et al., 2022). In particular, climate-mediated changes are predicted to continue altering marine food webs, with ecological, genetic, and economic consequences (Daufresne, Lengfellner, & Sommer, 2009; Pershing et al., 2015). Therefore, linking community-level processes to eco-evolutionary processes at the individual-level (such as polymorphic expression) is a prerequisite for understanding how taxa respond to these large-scale changes. Such an understanding is essential for effective conservation and ecologically sensitive fisheries management.

This study demonstrates that individual-level variation in prey acquisition both within and between sea-age phenotypes also displays a shared genetic basis (to varying degrees) with the *vgll3* locus. Furthermore, we show an apparent decrease in crustacean feeding efficiency for MSW salmon with the *vgll3*^EE^ genotype relative to 1SW salmon with the same genotype, but not for the *vgll3*^LL^ genotype. Taken together, these results are indicative of variation in feeding strategy acting as a functional ecological process that links *vgll3* genotype and sea-age in Atlantic salmon. Our results could help to provide a mechanism to explain the observed changes in the frequencies of 1SW and MSW phenotypes in Atlantic salmon populations (e.g., Czorlich, Aykanat, Erkinaro, Orell, & Primmer, 2018; Utne et al., 2021; Vollset et al., 2022). Changes in marine food webs have long been expected to negatively impact the energetic status of Atlantic salmon (Friedland, Chaput, & MacLean, 2005; B. Jonsson, Jonsson, & Albretsen, 2016; N. Jonsson & Jonsson, 2004; Strøm et al., 2023). The shared genetic basis between prey acquisition and sea-age indicates that Atlantic salmon sea-age is susceptible to evolution from human-induced changes to marine ecosystems.

## Acknowledgements

This study was funded by the Research Council of Finland (grants no. 328860, 353388 and 325964 to TA) and the Research Council of Norway (project number 280308 to KH). We acknowledge DNA Sequencing and Genomics Laboratory, Institute of Biotechnology, University of Helsinki for sequencing. We also thank Katja Enberg for providing comments to an earlier version of this manuscript.

## Author contributions

TA and KH conceptualized the paper. KH organized sample retrieval from scale archives at NINA. JAJ coordinated the Faroese fieldwork and collected the diet data. TA conducted the statistical analyses, genotyped the samples, and drafted the MS. All co-authors contributed to subsequent drafts.

## Data accessibility

The compiled datasets and R-codes to reproduce the results are available from the dryad repository https://doi.org/10.5061/dryad.fqz612k1m.

## Conflict of interest disclosure

We have no competing interests.

## Notes

### Competing Interest Statement

The authors have declared no competing interest.

### Summary of Updates

Changes have been made in parallel to the revisions of the version that is under peer-review.

## References

Amundsen, P. A., & Sanchez-Hernandez, J. (2019). Feeding studies take guts - critical review and recommendations of methods for stomach contents analysis in fish. Journal of Fish Biology. 95:1364–1373. doi.org/10.1111/jfb.14151

Aykanat, T., Jacobsen, J.A., Hindar, K. (2024). Diet drives performance trade-offs along the ontogeny in the Atlantic salmon. bioRxiv. doi: 10.1101/2024.06.04.597289

Aykanat, T., Jacobsen, J.A., Hindar, K. (2024) Ontogenetic variation in the marine foraging of Atlantic salmon functionally links genomic diversity with a major life history polymorphism. Dryad. doi.org/10.5061/dryad.fqz612k1m [Dataset]

Aykanat, T., Lindqvist, M., Pritchard, V. L., & Primmer, C. R. (2016). From population genomics to conservation and management: a workflow for targeted analysis of markers identified using genome-wide approaches in Atlantic salmon *Salmo salar*. Journal of Fish Biology, 89(6), 2658–2679. doi:10.1111/jfb.13149

Aykanat, T., Rasmussen, M., Ozerov, M., Niemela, E., Paulin, L., Vaha, J. P.,…Primmer, C. R. (2020). Life-history genomic regions explain differences in Atlantic salmon marine diet specialization. Journal of Animal Ecology, 89(11), 2677–2691. doi:10.1111/1365-2656.13324

Ayllon, F., Kjaerner-Semb, E., Furmanek, T., Wennevik, V., Solberg, M. F., Dahle, G.,... Wargelius, A. (2015). The vgll3 Locus Controls Age at Maturity in Wild and Domesticated Atlantic Salmon (*Salmo salar* L.) Males. PLoS Genetics, 11(11), e1005628. doi:10.1371/journal.pgen.1005628

Barson, N. J., Aykanat, T., Hindar, K., Baranski, M., Bolstad, G. H., Fiske, P.,…Primmer, C. R. (2015). Sex-dependent dominance at a single locus maintains variation in age at maturity in salmon. Nature, 528(7582), 405–408. doi:10.1038/nature16062

Bataillon, T., Gauthier, P., Villesen, P., Santoni, S., Thompson, J. D., & Ehlers, B. K. (2022). From genotype to phenotype: Genetic redundancy and the maintenance of an adaptive polymorphism in the context of high gene flow. Evolution Letters, 6(2), 189–202. doi:10.1002/evl3.277

Blasco-Moreno, A., M. Pérez-Casany, P. Puig, M. Morante, E. Castells, & O’Hara, R. B. (2019). What does a zero mean? Understanding false, random and structural zeros in ecology. Methods in Ecology and Evolution 10:949–959. doi.org/10.1111/2041-210X.13185

Bonnet, T., & Postma, E. (2018). Fluctuating selection and its (elusive) evolutionary consequences in a wild rodent population. Journal of Evolutionary Biology, 31(4), 572–586. doi:10.1111/jeb.13246

Brooks, M. E., Kristensen, K., van Benthem, K. J., Magnusson, A., Berg, C. W., Nielsen, A.,…Bolker, B. M. (2017). glmmTMB Balances Speed and Flexibility Among Packages for Zero-inflated Generalized Linear Mixed Modeling. R Journal, 9(2), 378–400. doi:Doi 10.32614/Rj-2017-066

Campbell, N. R., Harmon, S. A., & Narum, S. R. (2015). Genotyping-in-Thousands by sequencing (GT-seq): A cost effective SNP genotyping method based on custom amplicon sequencing. Molecular Ecology Resources, 15(4), 855–867. doi:10.1111/1755-0998.12357

Chaput, G., & Benoit, H. P. (2012). Evidence for bottom-up trophic effects on return rates to a second spawning for Atlantic salmon (Salmo salar) from the Miramichi River, Canada. ICES Journal of Marine Science, 69(9), 1656–1667. doi:10.1093/icesjms/fss055

Charmantier, A., Perrins, C., McCleery, R. H., & Sheldon, B. C. (2006). Quantitative genetics of age at reproduction in wild swans: Support for antagonistic pleiotropy models of senescence. Proceedings of National Academy of Sciences U S A, 103(17), 6587–6592. doi:10.1073/pnas.0511123103

Chouteau, M., Arias, M., & Joron, M. (2016). Warning signals are under positive frequency-dependent selection in nature. Proceedings of National Academy of Sciences U S A, 113(8), 2164–2169. doi:10.1073/pnas.1519216113

Coulson, T., MacNulty, D. R., Stahler, D. R., vonHoldt, B., Wayne, R. K., & Smith, D. W. (2011). Modeling effects of environmental change on wolf population dynamics, trait evolution, and life history. Science, 334(6060), 1275–1278. doi:10.1126/science.1209441

Cousminer, D. L., Berry, D. J., Timpson, N. J., Ang, W., Thiering, E., Byrne, E. M.,…Early Growth Genetics, C. (2013). Genome-wide association and longitudinal analyses reveal genetic loci linking pubertal height growth, pubertal timing and childhood adiposity. Human Molecular Genetics, 22(13), 2735–2747. doi:10.1093/hmg/ddt104

Czorlich, Y., Aykanat, T., Erkinaro, J., Orell, P., & Primmer, C. R. (2018). Rapid sex-specific evolution of age at maturity is shaped by genetic architecture in Atlantic salmon. Nature Ecology and Evolution, 2(11), 1800–1807. doi:10.1038/s41559-018-0681-5

Czorlich, Y., Aykanat, T., Erkinaro, J., Orell, P., & Primmer, C. R. (2022). Rapid evolution in salmon life history induced by direct and indirect effects of fishing. Science, 376(6591), 420–423. doi:10.1126/science.abg5980

Daufresne, M., Lengfellner, K., & Sommer, U. (2009). Global warming benefits the small in aquatic ecosystems. Proceedings of National Academy of Sciences U S A, 106(31), 12788–12793. doi:10.1073/pnas.0902080106

Debes, P. V., Piavchenko, N., Ruokolainen, A., Ovaskainen, O., Moustakas-Verho, J. E., Parre, N.,…Primmer, C. R. (2021). Polygenic and major-locus contributions to sexual maturation timing in Atlantic salmon. Molecular Ecology, 30(18), 4505–4519. doi:10.1111/mec.16062

Enberg, K., Jørgensen, C., Dunlop, E. S., Varpe, Ø., Boukal, D. S., Baulier, L.,…Heino, M. (2012). Fishing-induced evolution of growth: concepts, mechanisms and the empirical evidence. Marine Ecology, 33(1), 1–25. doi:10.1111/j.1439-0485.2011.00460.x

Esteve, M. (2005). Observations of Spawning Behaviour in Salmoninae: *Salmo*, *Oncorhynchus* and *Salvelinus*. Reviews in Fish Biology and Fisheries, 15(1-2), 1–21. doi:10.1007/s11160-005-7434-7

Ferguson, A., Reed, T. E., Cross, T. F., McGinnity, P., & Prodohl, P. A. (2019). Anadromy, potamodromy and residency in brown trout Salmo trutta: the role of genes and the environment. Journal of Fish Biology, 95(3), 692–718. doi:10.1111/jfb.14005

Fleming, I. A., & Einum, S. (2010). Reproductive Ecology: A Tale of Two Sexes. In Atlantic Salmon Ecology (pp. 33–65): Wiley-Blackwell.

Ford, E. B. (1945). Polymorphism. Biological Reviews, 20(2), 45–88. doi:10.1111/j.1469-185X.1945.tb00315.x

Friedland, K. D., Chaput, G., & MacLean, J. C. (2005). The emerging role of climate in post-smolt growth of Atlantic salmon. ICES Journal of Marine Science, 62(7), 1338–1349. doi:10.1016/j.icejms.2005.04.013

Funk, E. R., Mason, N. A., Palsson, S., Albrecht, T., Johnson, J. A., & Taylor, S. A. (2021). A supergene underlies linked variation in color and morphology in a Holarctic songbird. Nature Communications, 12(1), 6833. doi:10.1038/s41467-021-27173-z

Gilbey, J., Utne, K. R., Wennevik, V., Beck, A. C., Kausrud, K., Hindar, K.,…Verspoor, E. (2021). The early marine distribution of Atlantic salmon in the North-east Atlantic: A genetically informed stock-specific synthesis. Fish and Fisheries, 22(6), 1274–1306. doi:10.1111/faf.12587

Goldberg, J. K., Lively, C. M., Sternlieb, S. R., Pintel, G., Hare, J. D., Morrissey, M. B., & Delph, L. F. (2020). Herbivore-mediated negative frequency-dependent selection underlies a trichome dimorphism in nature. Evolution Letters, 4(1), 83–90. doi:10.1002/evl3.157

Grant, P. R., & Grant, B. R. (2002). Unpredictable evolution in a 30-year study of Darwin’s finches. Science, 296, 707–711.

Gross, M. R. (1985). Disruptive Selection for Alternative Life Histories in Salmon. Nature, 313(5997), 47–48. doi:Doi 10.1038/313047a0

Gross, M. R., Coleman, R. M., & McDowall, R. M. (1988). Aquatic productivity and the evolution of diadromous fish migration. Science, 239(4845), 1291–1293. doi:10.1126/science.239.4845.1291

Halperin, D. S., Pan, C., Lusis, A. J., & Tontonoz, P. (2013). Vestigial-like 3 is an inhibitor of adipocyte differentiation. Journal of Lipid Research, 54(2), 473–481. doi:10.1194/jlr.M032755

Hartig F (2022). DHARMa: Residual Diagnostics for Hierarchical (Multi-Level / Mixed) Regression Models. R package version 0.4.6.

Hedrick, P. W. (2012). What is the evidence for heterozygote advantage selection? Trends in Ecology & Evolution, 27(12), 698–704. doi:10.1016/j.tree.2012.08.012

Hvidsten, N. A., Jensen, A. J., Rikardsen, A. H., Finstad, B., Aure, J., Stefansson, S.,…Johnsen, B. O. (2009). Influence of sea temperature and initial marine feeding on survival of Atlantic salmon Salmo salar post-smolts from the Rivers Orkla and Hals, Norway. Journal of Fish Biology, 74(7), 1532–1548. doi:10.1111/j.1095-8649.2009.02219.x

Jacobsen, J. A., & Hansen, L. P. (2001). Feeding habits of wild and escaped farmed Atlantic salmon, *Salmo salar* L., in the Northeast Atlantic. ICES Journal of Marine Science, 58(4), 916–933. doi:10.1006/jmsc.2001.1084

Jacobsen, J. A., R. A. Lund, L. P. Hansen, & O’Maoileidigh, N. (2001). Seasonal differences in the origin of Atlantic salmon (Salmo salar L.) in the Norwegian Sea based on estimates from age structures and tag recaptures. Fisheries Research 52:169–177. doi.org/10.1016/S0165-7836(00)00255-1

Jamie, G. A., & Meier, J. I. (2020). The Persistence of Polymorphisms across Species Radiations. Trends in Ecology & Evolution, 35(9), 795–808. doi:10.1016/j.tree.2020.04.007

Janssen, K., Bustnes, J. O., & Mundy, N. I. (2021). Variation in Genetic Mechanisms for Plumage Polymorphism in Skuas (*Stercorarius*). Journal of Heredity, 112(5), 430–435. doi:10.1093/jhered/esab038

Johnston, S. E., Gratten, J., Berenos, C., Pilkington, J. G., Clutton-Brock, T. H., Pemberton, J. M., & Slate, J. (2013). Life history trade-offs at a single locus maintain sexually selected genetic variation. Nature, 502(7469), 93–95. doi:10.1038/nature12489

Jonsson, B., & Jonsson, N. (2011). Ecology of Atlantic salmon and brown trout : habitat as a template for life histories. Dordrecht: Springer.

Jonsson, B., Jonsson, N., & Albretsen, J. (2016). Environmental change influences the life history of salmon Salmo salar in the North Atlantic Ocean. Journal of Fish Biology, 88(2), 618–637. doi:10.1111/jfb.12854

Jonsson, N., & Jonsson, B. (2003). Energy allocation among developmental stages, age groups, and types of Atlantic salmon (Salmo salar) spawners. Canadian Journal of Fisheries and Aquatic Sciences, 60(5), 506–516. doi:10.1139/f03-042

Jonsson, N., & Jonsson, B. (2004). Size and age of maturity of Atlantic salmon correlate with the North Atlantic Oscillation Index (NAOI). Journal of Fish Biology, 64(1), 241–247. doi:10.1046/j.1095-8649.2004.00269.x

Keeley, E. R., & Grant, J. W. A. (1997). Allometry of diet selectivity in juvenile Atlantic salmon (*Salmo salar*). Canadian Journal of Fisheries and Aquatic Sciences, 54(8), 1894–1902. doi:10.1139/f97-096

Lee, B., Rizzoti, K., Kwon, D. S., Kim, S. Y., Oh, S., Epstein, D. J.,…Jeong, Y. (2012). Direct transcriptional regulation of Six6 is controlled by SoxB1 binding to a remote forebrain enhancer. Developmental Biology, 366(2), 393–403. doi:10.1016/j.ydbio.2012.04.023

Lenth R (2023). emmeans: Estimated Marginal Means, aka Least-Squares Means. R package version 1.8.5.

Marden, J. H., Langford, E. A., Robertson, M. A., & Fescemyer, H. W. (2021). Alleles in metabolic and oxygen-sensing genes are associated with antagonistic pleiotropic effects on life history traits and population fitness in an ecological model insect. Evolution, 75(1), 116–129. doi:10.1111/evo.14095

Martin, T. G., B. A. Wintle, J. R. Rhodes, P. M. Kuhnert, S. A. Field, S. J. Low-Choy, A. J. Tyre, & Possingham, H. P. (2005). Zero tolerance ecology: improving ecological inference by modelling the source of zero observations. Ecology Letters. 8:1235–1246. doi.org/10.1111/j.1461-0248.2005.00826.x

Matschiner, M., Barth, J. M. I., Torresen, O. K., Star, B., Baalsrud, H. T., Brieuc, M. S. O.,…Jentoft, S. (2022). Supergene origin and maintenance in Atlantic cod. Nature Ecology Evolution, 6(4), 469–481. doi:10.1038/s41559-022-01661-x

Mobley, K. B., Aykanat, T., Czorlich, Y., House, A., Kurko, J., Miettinen, A.,…Primmer, C. R. (2021). Maturation in Atlantic salmon (*Salmo salar*, Salmonidae): a synthesis of ecological, genetic, and molecular processes. Reviews in Fish Biology and Fisheries, 31(3), 523–571. doi:10.1007/s11160-021-09656-w

Mousseau, T. A., Sinervo, B., & Endler, J. A. (2000). Adaptive genetic variation in the wild. New York: Oxford University Press.

Moustakas-Verho, J. E., Kurko, J., House, A. H., Erkinaro, J., Debes, P., & Primmer, C. R. (2020). Developmental expression patterns of six6: A gene linked with spawning ecotypes in Atlantic salmon. Gene Expr Patterns, 38, 119149. doi:10.1016/j.gep.2020.119149

O’Sullivan, R. J., Ozerov, M., Bolstad, G. H., Gilbey, J., Jacobsen, J. A., Erkinaro, J.,…Grant, W. S. (2022). Genetic stock identification reveals greater use of an oceanic feeding ground around the Faroe Islands by multi-sea winter Atlantic salmon, with variation in use across reporting groups. ICES Journal of Marine Science, 79(9), 2442–2452. doi:10.1093/icesjms/fsac182

Pershing, A. J., Alexander, M. A., Hernandez, C. M., Kerr, L. A., Le Bris, A., Mills, K. E.,…Thomas, A. C. (2015). Slow adaptation in the face of rapid warming leads to collapse of the Gulf of Maine cod fishery. Science, 350(6262), 809–812. doi:10.1126/science.aac9819

R Core Team. (2019). R: A language and environment for statistical computing. In: R Foundation for Statistical Computing, Vienna, Austria.

Reznick, D. (2016). Hard and Soft Selection Revisited: How Evolution by Natural Selection Works in the Real World. Journal of Heredity, 107(1), 3–14. doi:10.1093/jhered/esv076

Rikardsen, A. H., & Dempson, J. B. (2010). Dietary Life-Support: The Food and Feeding of Atlantic Salmon at Sea. In Atlantic Salmon Ecology (pp. 115–143): Wiley-Blackwell.

Salminen, M., Erkamo, E., & Salmi, J. (2001). Diet of post-smolt and one-sea-winter Atlantic salmon in the Bothnian Sea, Northern Baltic. Journal of Fish Biology, 58(1), 16–35. doi:DOI 10.1006/jfbi.2000.1426

Salminen, M., Kuikka, S., & Erkamo, E. (1995). Annual variability in survival of sea-ranched Baltic salmon,Salmo salarL: significance of smolt size and marine conditions. Fisheries Management and Ecology, 2(3), 171–184. doi:10.1111/j.1365-2400.1995.tb00110.x

Sinclair-Waters, M., Piavchenko, N., Ruokolainen, A., Aykanat, T., Erkinaro, J., & Primmer, C. R. (2022). Refining the genomic location of single nucleotide polymorphism variation affecting Atlantic salmon maturation timing at a key large-effect locus. Molecular Ecology, 31(2), 562–570. doi:10.1111/mec.16256

Strøm, J. F., Ugedal, O., Rikardsen, A. H., & Thorstad, E. B. (2023). Marine food consumption by adult Atlantic salmon and energetic impacts of increased ocean temperatures caused by climate change. Hydrobiologia, 850(14), 3077–3089. doi:10.1007/s10750-023-05234-2

Svensson, E. I., Willink, B., Duryea, M. C., & Lancaster, L. T. (2020). Temperature drives pre-reproductive selection and shapes the biogeography of a female polymorphism. Ecology Letters, 23(1), 149–159. doi:10.1111/ele.13417

Sydeman, W. J., Poloczanska, E., Reed, T. E., & Thompson, S. A. (2015). Climate change and marine vertebrates. Science, 350(6262), 772–777. doi:10.1126/science.aac9874

Thorpe, J. E., Mangel, M., Metcalfe, N. B., & Huntingford, F. A. (1998). Modelling the proximate basis of salmonid life-history variation, with application to Atlantic salmon, *Salmo salar* L. Evolutionary Ecology, 12(5), 581–599. doi:Doi 10.1023/A:1022351814644

Utne, K. R., Pauli, B. D., Haugland, M., Jacobsen, J. A., Maoileidigh, N., Melle, W.,…Secor, D. (2021). Poor feeding opportunities and reduced condition factor for salmon post-smolts in the Northeast Atlantic Ocean. ICES Journal of Marine Science, 78(8), 2844–2857. doi:10.1093/icesjms/fsab163

Vollset, K. W., Urdal, K., Utne, K., Thorstad, E. B., Saegrov, H., Raunsgard, A.,…Fiske, P. (2022). Ecological regime shift in the Northeast Atlantic Ocean revealed from the unprecedented reduction in marine growth of Atlantic salmon. Science Advances, 8(9), eabk2542. doi:10.1126/sciadv.abk2542

Youngson, A. F., & McLay, H. A. (1985). Changes in steroid hormone levels in blood serum as an early predictor of approaching maturity in Atlantic salmon. ICES Journal of Marine Science, 69(1-2), 145–157. doi.org/10.1016/0044-8486(88)90193-7

Wańkowski, J. W. J., & Thorpe, J. E. (2006). Spatial distribution and feeding in atlantic salmon, Salmo salar L. juveniles. Journal of Fish Biology, 14(3), 239–247. doi:10.1111/j.1095-8649.1979.tb03515.x

