## Supplementary Materials for "Ontogenetic variation in the marine foraging of Atlantic salmon functionally links genomic diversity with a major life history polymorphism"

for

### Supplementary Tables

**Supplementary Table 1:** The details of diet in Atlantic salmon sub-adult stomachs used in the study (N=1536).

| Diet type | Species | Taxonomic groups | Number if stomachs with diet | Total weight (grams) | Total number of item | Average weight (grams) |
|---|---|---|---|---|---|---|
| **Crustaceans** | *Meganyctiphanes norvegica* | Euphausiids | 626 | 221.66 | 1941 | 0.114 |
| | *Themisto libellula* | Hyperiid amphipods | 407 | 198.869 | 2024 | 0.098 |
| | *Themisto compressa* | Hyperiid amphipods | 360 | 659.817 | 24667 | 0.027 |
| | *Hymenodora glacialis* | shrimp | 247 | 244 | 342 | 0.713 |
| | *Themisto spp* | Hyperiid amphipods | 206 | 241.418 | 6480 | 0.037 |
| | *Themisto compressa f. bispinosa* | Hyperiid amphipods | 83 | 43.375 | 2058 | 0.021 |
| | *Themisto compressa f. compressa* | Hyperiid amphipods | 53 | 23.036 | 867 | 0.027 |
| | *Ephausiidae* | Euphausiids | 44 | 25.04 | 305 | 0.082 |
| | *Themisto abyssorum* | Hyperiid amphipods | 27 | 1.86 | 33 | 0.056 |
| | *Eusirus holmi* | amphipod | 26 | 14.12 | 27 | 0.523 |
| | *Thysanoessa inermis* | Euphausiids | 17 | 3.83 | 44 | 0.087 |
| | *Sergestes arcticus* | shrimp | 7 | 5.36 | 7 | 0.766 |
| | Gammaridea | | 3 | 0.93 | 3 | 0.310 |
| | *Pasiphea tarda* | shrimp | 1 | 0.89 | 1 | 0.890 |
| | *Thysanoessa longicaudata* | Euphausiids | 1 | 0.004 | 1 | 0.004 |
| **Fish** | *Maurolicus muelleri* | Pearlsides | 324 | 960.97 | 866 | 1.110 |
| | *Benthosema glaciale* | Lanternfishes | 268 | 811.632 | 375 | 2.164 |
| | Myctophidae | Lanternfishes | 117 | 617.9 | 293 | 2.109 |
| | Fry/larvae | mostly *Mallotus villosus* | 66 | 111.76 | 424 | 0.264 |
| | Paralepidae | Baracudinas | 35 | 749.78 | 37 | 20.264 |
| | *Myctophum punctatum* | Lanternfishes | 16 | 44.67 | 23 | 1.942 |
| | *Micromesistius poutassou* | blue whiting | 10 | 390.39 | 10 | 39.039 |
| | *Notolepis rissoi kroyeri* | Baracudinas | 9 | 189.64 | 9 | 21.071 |
| | *Clupea harengus* | Herring | 5 | 360.23 | 5 | 72.046 |
| | *Notoscopelus kroeyeri* | Lanternfishes | 5 | 41.28 | 6 | 6.880 |
| | *Scomber scombrus* | Mackerel | 4 | 151.95 | 4 | 37.988 |
| | Ammodyteidae | Sandlance | 3 | 2.71 | 4 | 0.678 |
| | *Gasterosteus aculeatus* | Three-spined stickleback | 3 | 8.5 | 3 | 2.833 |
| | *Mallotus villosus* | Capelin | 2 | 40.96 | 3 | 13.653 |
| | *Paralepis coregonoides borealis* | Baracudinas | 2 | 62.13 | 2 | 31.065 |
| | *Lampanyctus crocodilus* | Lanternfishes | 1 | 13.52 | 1 | 13.520 |
| | *Lycenchelys spp* | Eelpouts | 1 | 8 | 1 | 8.000 |
| **Others**[1] | Gonatidae | Squid | 11 | 180.39 | 8[1] | NA |
| | Organic remains, | | 17 | 68.24 | NA | NA |
| | Crustacean remains | | 255 | 263.99 | 13[1] | NA |
| | Fish remains | | 527 | 1648.59 | 22[1] | NA |

[1] Only a subset of remains were enumerated as individual items. Squids, and if enumeration was possible, fish remains were counted within the fish group, and crustaceans remains with the crustaceans group.

**Supplementary Table 2:** Presence and absence of crustaceans and fish diets in the stomach of Atlantic salmon as a function of sea-age.

| | 1SW | 2SW | 3SW |
|---|---|---|---|
| **Crustaceans present in the diet** | 68 | 526 | 234 |
| **No crustaceans present in the diet** | 100 | 424 | 184 |
| **Fish present in the diet** | 23 | 289 | 151 |
| **No fish present in the diet** | 145 | 661 | 267 |

**Supplementary Table 3:** Fixed coefficients of the zero-inflated model with a negative binomial error distribution in the count component to quantify crustacean diet in the stomach of sub-adult Atlantic salmon

| | Parameter | Estimate | Std. Error | *z* value | *p* value |
|---|---|---|---|---|---|
| **Count component** | Intercept | 4.364 | 0.442 | 9.866 | < 0.001 |
| | Season (winter) | -1.777 | 0.348 | -5.107 | < 0.001 |
| | Fishing period (93/94) | 1.591 | 0.289 | 5.512 | < 0.001 |
| | Regional group (Northern Norway) | -0.480 | 0.192 | -2.504 | 0.012 |
| | Regional group (Southern Norway) | -0.290 | 0.156 | -1.858 | 0.063 |
| | Sea age | -0.512 | 0.114 | -4.478 | < 0.001 |
| | $L_{age.std}$ (scaled) | 0.142 | 0.211 | 0.675 | 0.500 |
| | *Vgll3* (scaled) | 0.033 | 0.230 | 0.142 | 0.887 |
| | *Six6* (scaled) | 0.136 | 0.225 | 0.606 | 0.544 |
| | Sex (scaled) | 0.151 | 0.205 | 0.736 | 0.462 |
| | Sea age : $L_{age.std}$ (scaled) | -0.100 | 0.089 | -1.122 | 0.262 |
| | Sea age : *Vgll3* (scaled) | 0.033 | 0.103 | 0.320 | 0.749 |
| | Sea age : *Six6* (scaled) | -0.026 | 0.105 | -0.248 | 0.804 |
| | Sea age : Sex (scaled) | -0.063 | 0.090 | -0.703 | 0.482 |
| **Zero inflation component** | Intercept | 1.326 | 0.703 | 1.887 | 0.059 |
| | Season (winter) | -1.133 | 0.464 | -2.439 | 0.015 |
| | Fishing period (93/94) | 0.039 | 0.427 | 0.092 | 0.927 |
| | Regional group (Northern Norway) | -1.627 | 0.682 | -2.387 | 0.017 |
| | Regional group (Southern Norway) | -0.330 | 0.271 | -1.217 | 0.224 |
| | Sea age | -0.861 | 0.270 | -3.196 | 0.001 |
| | $L_{age.std}$ (scaled) | -0.140 | 0.332 | -0.422 | 0.673 |
| | *Vgll3* (scaled) | -0.945 | 0.428 | -2.208 | 0.027 |
| | *Six6* (scaled) | 0.574 | 0.383 | 1.499 | 0.134 |
| | Sex (scaled) | 1.133 | 0.378 | 2.998 | 0.003 |
| | Sea age : $L_{age.std}$ (scaled) | 0.044 | 0.171 | 0.254 | 0.799 |
| | Sea age : *Vgll3* (scaled) | 0.612 | 0.228 | 2.690 | 0.007 |
| | Sea age : *Six6* (scaled) | -0.363 | 0.205 | -1.769 | 0.077 |
| | Sea age : Sex (scaled) | -0.576 | 0.204 | -2.822 | 0.005 |

**Supplementary Table 4:** Fixed coefficients of the zero-inflated model with a negative binomial error distribution in the count component to quantify Hyperiid amphipods (mostly *Themisto* species) diet in the stomach of sub-adult Atlantic salmon.

| | Parameter | Estimate | Std. Error | *z* value | *p* value |
|---|---|---|---|---|---|
| **Count component** | Intercept | 4.795 | 0.536 | 8.948 | < 0.001 |
| | Season (winter) | -2.217 | 0.415 | -5.340 | < 0.001 |
| | Fishing period (93/94) | 1.643 | 0.357 | 4.603 | < 0.001 |
| | Regional group (Northern Norway) | -0.594 | 0.239 | -2.485 | 0.013 |
| | Regional group (Southern Norway) | -0.416 | 0.191 | -2.179 | 0.029 |
| | Sea age | -0.738 | 0.144 | -5.108 | < 0.001 |
| | $L_{age.std}$ (scaled) | 0.307 | 0.259 | 1.187 | 0.235 |
| | *Vgll3* (scaled) | -0.059 | 0.287 | -0.207 | 0.836 |
| | *Six6* (scaled) | 0.263 | 0.270 | 0.973 | 0.330 |
| | Sex (scaled) | 0.281 | 0.254 | 1.107 | 0.268 |
| | Sea age : $L_{age.std}$ (scaled) | -0.198 | 0.108 | -1.838 | 0.066 |
| | Sea age : *Vgll3* (scaled) | 0.097 | 0.131 | 0.741 | 0.459 |
| | Sea age : *Six6* (scaled) | -0.078 | 0.127 | -0.615 | 0.538 |
| | Sea age : Sex (scaled) | -0.129 | 0.112 | -1.149 | 0.250 |
| **Zero inflation component** | Intercept | 1.389 | 0.750 | 1.852 | 0.064 |
| | Season (winter) | -0.847 | 0.531 | -1.595 | 0.111 |
| | Fishing period (93/94) | -0.470 | 0.482 | -0.974 | 0.330 |
| | Regional group (Northern Norway) | -1.047 | 0.552 | -1.898 | 0.058 |
| | Regional group (Southern Norway) | -0.259 | 0.287 | -0.900 | 0.368 |
| | Sea age | -0.731 | 0.271 | -2.701 | 0.007 |
| | $L_{age.std}$ (scaled) | -0.188 | 0.345 | -0.544 | 0.586 |
| | *Vgll3* (scaled) | -0.703 | 0.440 | -1.597 | 0.110 |
| | *Six6* (scaled) | 0.500 | 0.395 | 1.268 | 0.205 |
| | Sex (scaled) | 1.130 | 0.391 | 2.892 | 0.004 |
| | Sea age : $L_{age.std}$ (scaled) | 0.077 | 0.173 | 0.445 | 0.656 |
| | Sea age : *Vgll3* (scaled) | 0.482 | 0.234 | 2.060 | 0.039 |
| | Sea age : *Six6* (scaled) | -0.272 | 0.210 | -1.294 | 0.196 |
| | Sea age : Sex (scaled) | -0.555 | 0.209 | -2.655 | 0.008 |

**Supplementary Table 5:** Fixed coefficients of the zero-inflated model with a negative binomial error distribution in the count component to quantify Euphausiids (krill) diet in the stomach of sub-adult Atlantic salmon.

| | Parameter | Estimate | Std. Error | *z* value | *p* value |
|---|---|---|---|---|---|
| **Count component** | Intercept | -0.469 | 0.480 | -0.978 | 0.328 |
| | Season (winter) | 1.411 | 0.371 | 3.806 | < 0.001 |
| | Fishing period (93/94) | 0.508 | 0.298 | 1.705 | 0.088 |
| | Regional group (Northern Norway) | -0.280 | 0.240 | -1.164 | 0.245 |
| | Regional group (Southern Norway) | -0.027 | 0.209 | -0.131 | 0.896 |
| | Sea age | -0.168 | 0.139 | -1.206 | 0.228 |
| | $L_{age.std}$ (scaled) | 0.186 | 0.368 | 0.505 | 0.614 |
| | *Vgll3* (scaled) | 0.212 | 0.317 | 0.670 | 0.503 |
| | *Six6* (scaled) | -0.157 | 0.331 | -0.474 | 0.636 |
| | Sex (scaled) | -0.256 | 0.278 | -0.920 | 0.358 |
| | Sea age : $L_{age.std}$ (scaled) | -0.080 | 0.160 | -0.498 | 0.619 |
| | Sea age : *Vgll3* (scaled) | -0.041 | 0.138 | -0.297 | 0.766 |
| | Sea age : *Six6* (scaled) | 0.009 | 0.149 | 0.061 | 0.952 |
| | Sea age : Sex (scaled) | 0.127 | 0.124 | 1.018 | 0.308 |
| **Zero inflation component** | Intercept | 0.710 | 0.799 | 0.889 | 0.374 |
| | Season (winter) | 0.016 | 0.561 | 0.028 | 0.977 |
| | Fishing period (93/94) | -0.240 | 0.379 | -0.633 | 0.527 |
| | Regional group (Northern Norway) | -1.147 | 0.648 | -1.771 | 0.077 |
| | Regional group (Southern Norway) | -0.324 | 0.352 | -0.920 | 0.357 |
| | Sea age | -0.421 | 0.283 | -1.488 | 0.137 |
| | $L_{age.std}$ (scaled) | -0.275 | 0.624 | -0.440 | 0.660 |
| | *Vgll3* (scaled) | -0.790 | 0.534 | -1.481 | 0.139 |
| | *Six6* (scaled) | -0.161 | 0.502 | -0.320 | 0.749 |
| | Sex (scaled) | 0.144 | 0.448 | 0.321 | 0.748 |
| | Sea age : $L_{age.std}$ (scaled) | 0.130 | 0.287 | 0.454 | 0.650 |
| | Sea age : *Vgll3* (scaled) | 0.418 | 0.250 | 1.671 | 0.095 |
| | Sea age : *Six6* (scaled) | -0.043 | 0.242 | -0.177 | 0.860 |
| | Sea age : Sex (scaled) | -0.085 | 0.214 | -0.398 | 0.691 |

**Supplementary Table 6:** The likelihood of zero-inflated negative binomial models with alternative parameterization of genotype (*vgll3* and *six6*) and sea-age.

| Genotype and sea-age parameterizations | | | AIC | |
|---|---|---|---|---|
| ***vgll3*** | ***six6*** | **sea age** | **Crustaceans model** | **Fish models** |
| additive | additive | numeric | 7986.44 | 3606.19 |
| non-additive | non-additive | numeric | 7996.56 | 3609.98 |
| non-additive | additive | numeric | 7991.61 | 3608.31 |
| additive | non-additive | numeric | 7990.95 | 3606.56 |
| additive | additive | categorical | 7992.16 | No convergence |
| non-additive | non-additive | categorical | 8008.48 | No convergence |
| non-additive | additive | categorical | 8000.19 | No convergence |
| additive | non-additive | categorical | 7998.77 | No convergence |

**Supplementary Table 7:** The likelihood of zero-inflated negative binomial models with alternative interaction models between sea-age, sex and, genotype (*vgll3* and *six6*).

| Model structure | Definition | AIC | |
|---|---|---|---|
| | | **Crustaceans model** | **Fish models** |
| sea age * (*vgll3 + six6 + sex*) | Default model | 7986.44 | 3606.19 |
| sea age * (*vgll3 *sex + six6*) | Three way interaction with sex, sea-age and genotype (*vgll3*) | 7989.05 | 3608.27 |
| sea age * (*vgll3 + six6 * sex*) | Three way interaction with sex, sea-age and genotype (*six6*) | 7987.33 | 3608.99 |
| sea age * sex * (*vgll3 + six6*) | Three way interaction with sex, sea-age and genotype (*six6* and *vgll3* separately) | 7990.10 | 3612.52 |
| sea age * sex * (*vgll3 * six6*) | Four way interaction with sex, sea-age and genotype (*six6* and *vgll3*) | 7997.09 | No convergence |

**Supplementary Table 8:** Fixed coefficients of the zero-inflated model with a negative binomial error distribution in the count component to quantify fish diet in the stomach of sub-adult Atlantic salmon

| | Parameter | Estimate | Std. Error | *z* value | *p* value |
|---|---|---|---|---|---|
| **Count component** | Intercept | -0.279 | 0.781 | -0.358 | 0.721 |
| | Season (winter) | 2.631 | 0.596 | 4.413 | < 0.001 |
| | Fishing period (93/94) | 0.141 | 0.331 | 0.425 | 0.671 |
| | Regional group (Northern Norway) | -0.225 | 0.265 | -0.849 | 0.396 |
| | Regional group (Southern Norway) | -0.162 | 0.228 | -0.711 | 0.477 |
| | Sea age | -0.738 | 0.175 | -4.204 | < 0.001 |
| | $L_{age.std}$ (scaled) | -0.613 | 0.396 | -1.547 | 0.122 |
| | *Vgll3* (scaled) | 0.754 | 0.361 | 2.088 | 0.037 |
| | *Six6* (scaled) | 0.334 | 0.343 | 0.974 | 0.330 |
| | Sex (scaled) | -0.821 | 0.340 | -2.416 | 0.016 |
| | Sea age : $L_{age.std}$ (scaled) | 0.137 | 0.170 | 0.805 | 0.421 |
| | Sea age : *Vgll3* (scaled) | -0.278 | 0.155 | -1.796 | 0.072 |
| | Sea age : *Six6* (scaled) | -0.219 | 0.156 | -1.403 | 0.160 |
| | Sea age : Sex (scaled) | 0.366 | 0.147 | 2.484 | 0.013 |
| **Zero inflation component** | Intercept | 6.353 | 1.737 | 3.658 | < 0.001 |
| | Season (winter) | -1.962 | 0.974 | -2.014 | 0.044 |
| | Fishing period (93/94) | -1.593 | 0.695 | -2.293 | 0.022 |
| | Regional group (Northern Norway) | -0.949 | 0.667 | -1.423 | 0.155 |
| | Regional group (Southern Norway) | -0.814 | 0.625 | -1.303 | 0.193 |
| | Sea age | -2.444 | 0.660 | -3.704 | < 0.001 |
| | $L_{age.std}$ (scaled) | -1.044 | 1.280 | -0.816 | 0.415 |
| | *Vgll3* (scaled) | -0.434 | 0.899 | -0.483 | 0.629 |
| | *Six6* (scaled) | 0.789 | 0.896 | 0.881 | 0.379 |
| | Sex (scaled) | -2.000 | 1.039 | -1.925 | 0.054 |
| | Sea age : $L_{age.std}$ (scaled) | 0.049 | 0.552 | 0.088 | 0.930 |
| | Sea age : *Vgll3* (scaled) | 0.414 | 0.473 | 0.875 | 0.382 |
| | Sea age : *Six6* (scaled) | -0.349 | 0.467 | -0.749 | 0.454 |
| | Sea age : Sex (scaled) | 1.180 | 0.555 | 2.127 | 0.033 |

**Supplementary Table 9:** Fish model parameters with addition of dispersion parameters for ill-fitted variables, as identified by diagnostic plots.

| | Parameter | Estimate | Std. Error | *z* value | *p* value |
|---|---|---|---|---|---|
| **Count component** | Intercept | 0.040 | 0.772 | 0.052 | 0.959 |
| | Season (winter) | 2.048 | 0.697 | 2.936 | 0.003 |
| | Fishing period (93/94) | 0.346 | 0.332 | 1.042 | 0.297 |
| | Regional group (Northern Norway) | -0.113 | 0.257 | -0.441 | 0.659 |
| | Regional group (Southern Norway) | -0.070 | 0.250 | -0.279 | 0.780 |
| | Sea age | -0.723 | 0.159 | -4.533 | < 0.001 |
| | $L_{age.std}$ (scaled) | -0.574 | 0.442 | -1.301 | 0.193 |
| | *Vgll3* (scaled) | 0.799 | 0.348 | 2.293 | 0.022 |
| | *Six6* (scaled) | 0.272 | 0.347 | 0.785 | 0.433 |
| | Sex (scaled) | -0.704 | 0.334 | -2.106 | 0.035 |
| | Sea age : $L_{age.std}$ (scaled) | 0.138 | 0.185 | 0.747 | 0.455 |
| | Sea age : *Vgll3* (scaled) | -0.274 | 0.154 | -1.780 | 0.075 |
| | Sea age : *Six6* (scaled) | -0.159 | 0.162 | -0.986 | 0.324 |
| | Sea age : Sex (scaled) | 0.301 | 0.149 | 2.016 | 0.044 |
| **Zero inflation component** | Intercept | 6.760 | 1.838 | 3.678 | < 0.001 |
| | Season (winter) | -3.460 | 1.273 | -2.719 | 0.007 |
| | Fishing period (93/94) | -0.481 | 0.788 | -0.611 | 0.542 |
| | Regional group (Northern Norway) | 0.472 | 0.799 | 0.591 | 0.554 |
| | Regional group (Southern Norway) | -0.268 | 0.777 | -0.345 | 0.730 |
| | Sea age | -2.647 | 0.779 | -3.399 | 0.001 |
| | $L_{age.std}$ (scaled) | -0.694 | 1.253 | -0.554 | 0.579 |
| | *Vgll3* (scaled) | -0.778 | 1.229 | -0.633 | 0.527 |
| | *Six6* (scaled) | 0.649 | 1.052 | 0.617 | 0.537 |
| | Sex (scaled) | -1.560 | 1.084 | -1.439 | 0.150 |
| | Sea age : $L_{age.std}$ (scaled) | -0.134 | 0.568 | -0.236 | 0.814 |
| | Sea age : *Vgll3* (scaled) | 0.737 | 0.777 | 0.949 | 0.343 |
| | Sea age : *Six6* (scaled) | -0.202 | 0.606 | -0.334 | 0.739 |
| | Sea age : Sex (scaled) | 0.902 | 0.644 | 1.400 | 0.162 |
| **Dispersion model** | Intercept | -0.081 | 2.173 | -0.037 | 0.970 |
| | Season (winter) | -1.655 | 2.200 | -0.752 | 0.452 |
| | Regional group (Northern Norway) | 1.156 | 0.308 | 3.748 | < 0.001 |
| | Regional group (Southern Norway) | 0.385 | 0.229 | 1.686 | 0.092 |
| | Fishing period (93/94) | 0.674 | 0.228 | 2.955 | 0.003 |
| | Sex (scaled) | -0.028 | 0.282 | -0.098 | 0.922 |

**Supplementary Table 10:** Effect of vgll3 and six6 genotype to condition.

| Parameter | Estimate | Std. Error | *z* value | *p* value |
|---|---|---|---|---|
| Intercept | 1.081 | 0.018 | 61.311 | < 0.001 |
| Season (winter) | -0.082 | 0.012 | -6.713 | < 0.001 |
| Fishing period (93/94) | -0.019 | 0.010 | -1.889 | 0.059 |
| Regional group (Northern Norway) | -0.052 | 0.010 | -4.985 | < 0.001 |
| Regional group (Southern Norway) | -0.048 | 0.008 | -6.042 | < 0.001 |
| Sea age (SW2) | 0.034 | 0.013 | 2.645 | 0.008 |
| Sea age (SW3) | 0.182 | 0.015 | 12.217 | < 0.001 |
| $L_{age.std}$ (scaled) | -0.009 | 0.008 | -1.144 | 0.253 |
| *Vgll3* (scaled) | 0.006 | 0.010 | 0.651 | 0.515 |
| *Six6* (scaled) | 0.013 | 0.008 | 1.511 | 0.131 |
| Sex (scaled) | 0.016 | 0.008 | 1.982 | 0.047 |
| Sea age (SW2):$L_{age.std}$ (scaled) | 0.031 | 0.008 | 3.661 | < 0.001 |
| Sea age (SW3):$L_{age.std}$ (scaled) | 0.021 | 0.010 | 2.080 | 0.038 |
| Sea age (SW2):*Vgll3* (scaled) | -0.008 | 0.010 | -0.759 | 0.448 |
| Sea age (SW3):*Vgll3* (scaled) | -0.004 | 0.012 | -0.315 | 0.753 |
| Sea age (SW2):*Six6* (scaled) | -0.036 | 0.009 | -3.972 | < 0.001 |
| Sea age (SW3):*Six6* (scaled) | -0.009 | 0.012 | -0.741 | 0.459 |
| Sea age (SW2):Sex (scaled) | -0.034 | 0.009 | -3.787 | < 0.001 |
| Sea age (SW3):Sex (scaled) | -0.018 | 0.010 | -1.773 | 0.076 |

### Supplementary Figures

**Supplementary Figure 1:** Distribution (numbers) of crustaceans (a) and fish (b) diet in the stomachs of Atlantic salmon in this study (n=1536).

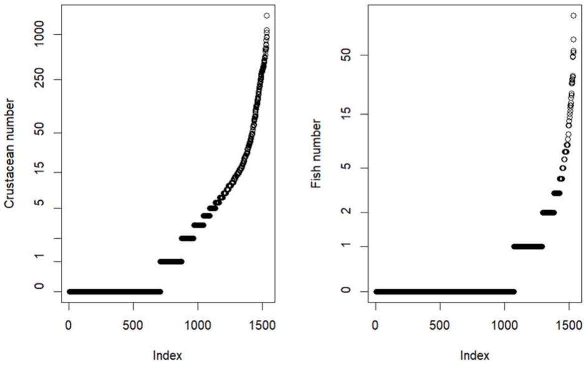

**Supplementary Figure 2:** Diagnostic plots for the crustaceans model, the fish model, and the fish model after dispersion parameters were included.

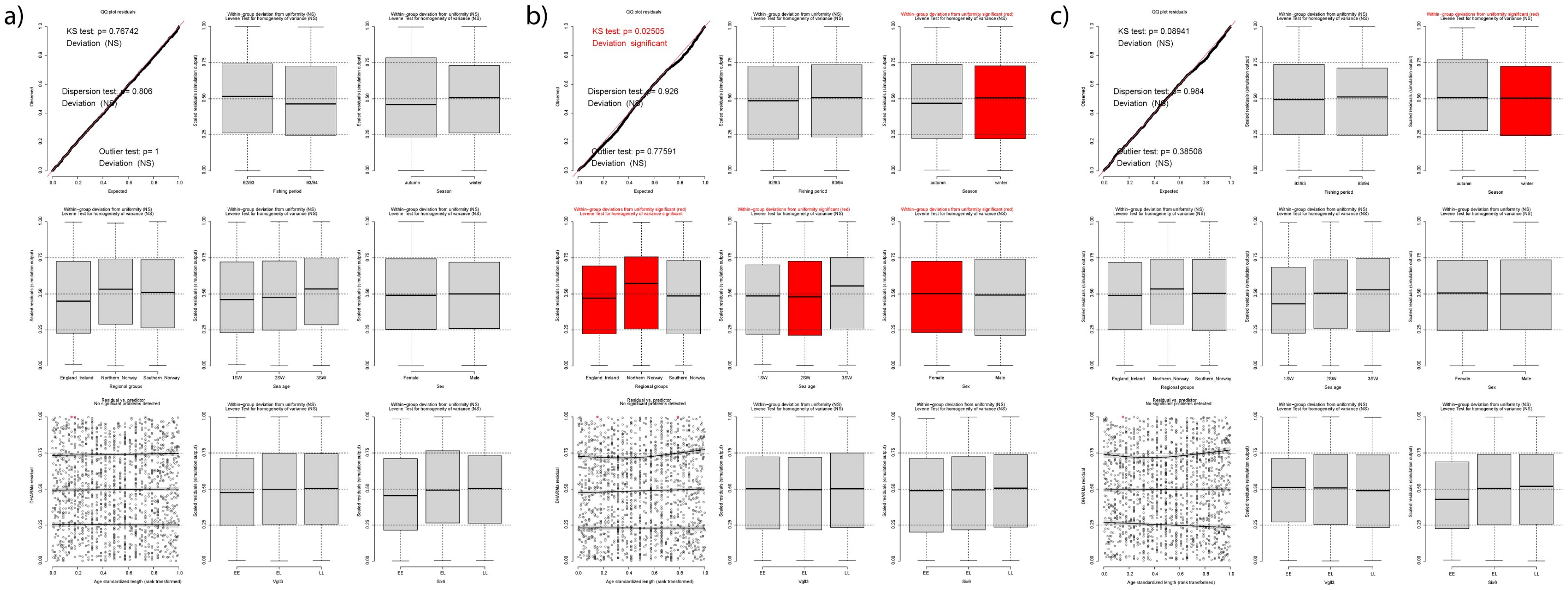

**Supplementary Figure 3:** Marginal *vgll3* effect to foraging outcome (count component) in the fish model when age-standardized length ($L_{age.std}$ ) was excluded from the model.

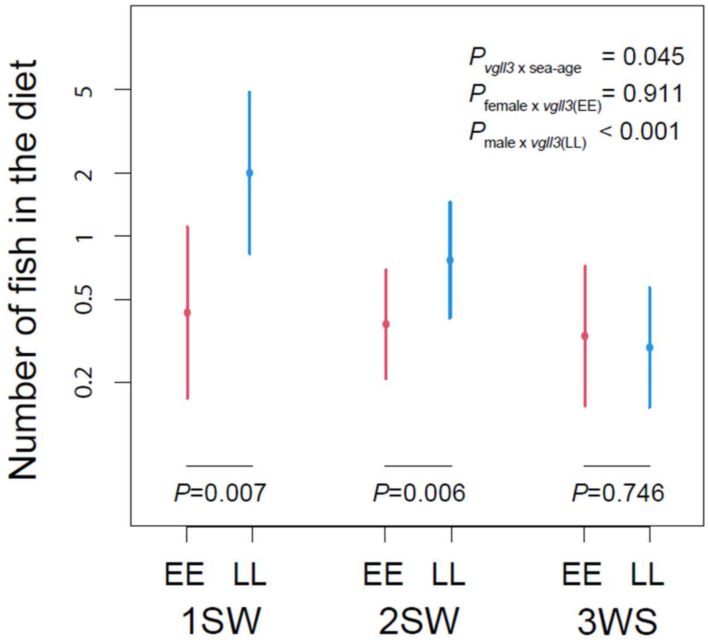
